# Lactylation-driven FTO-mediated m^6^A modification of CDK2 aggravates diabetic microvascular anomalies

**DOI:** 10.1101/2023.05.19.541470

**Authors:** Xue Chen, Ru-Xu Sun, Jia-Nan Wang, Ye-Ran Zhang, Bing Qin, Yi-Chen Zhang, Yuan-Xin Dai, Hong-Jing Zhu, Ying Wang, Jin-Xiang Zhao, Wei-Wei Zhang, Jiang-Dong Ji, Song-Tao Yuan, Qun-Dong Shen, Qing-Huai Liu

## Abstract

Diabetic retinopathy (DR) is a leading cause of irreversible vision loss in working-age populations. FTO is an N^6^-methyladenosine (m^6^A) demethylase that participates in various biological events, while its role in DR remains elusive. Herein, we detected elevated FTO expression in retinal proliferative membranes of DR patients. FTO promoted endothelial cell (EC) cell cycle progression and tip cell formation to facilitate angiogenesis *in vitro*, in mice and in zebrafish. FTO also regulated EC-pericyte crosstalk to trigger diabetic microvascular leakage, and mediated EC-microglia interactions to induce retinal inflammation and neurodegeneration *in vivo* and *in vitro*. Mechanistically, FTO affected EC features via modulating *CDK2* mRNA stability in an m^6^A-YTHDF2-dependent manner. FTO up-regulation under diabetic conditions was driven by lactate mediated histone lactylation. FB23-2, an inhibitor to FTO’s m^6^A demethylase activity, suppressed angiogenic phenotypes *in vivo* and *in vitro*. Noteworthy, we developed a nanoplatform encapsulating FB23-2 for systemic administration, and confirmed its targeting and therapeutic efficiencies in mice. Collectively, our study demonstrated that FTO coordinates EC biology and retinal homeostasis in DR, providing a promising nanotherapeutic approach for DR.

## Introduction

Diabetic retinopathy (DR), a major microvascular complication of diabetes, is emerging as a leading threat to vision in working-age populations. Clinically, DR is divided into the early and the advanced stages based on its disease course. Non-proliferative DR (NPDR) represents the early stage of DR, which is characterized by increased vascular permeability and capillary occlusions. Proliferative DR (PDR) is the advanced form of DR with the clinical hallmark of neovascularization. Patients may be asymptomatic in NPDR but may experience severe vision loss in PDR when vitreous hemorrhage or tractional retinal detachment happens. Retinal microvasculopathy, inflammation and neurodegeneration are major pathological features of DR (1). Microvascular endothelial cells (ECs) are major targets of hyperglycemic injuries and are mostly studied in DR (2). Loss of cell-to-cell contact in EC monolayers contributes to increased blood-retinal barrier (BRB) permeability, and its transformation into tip cells leads to sprouting angiogenesis (3). Pericyte is another major cellular constituent in neural retinal microvessels. Formation, maturation, and stabilization of the micro-vasculatures require EC-pericyte interactions, which are perturbed in DR, resulting in BRB rupture and other microangiopathies (4). Interrupted homeostasis in diabetic retina induces microglia activation and inflammatory responses, which drive sustained vascular damages, further resulting in increased vascular permeability and angiogenesis [3,4]. Retinal neurodegeneration, especially axonal degeneration of retinal ganglion cell (RGC), is one of the earliest events in DR progression (3). Loss of RGCs, the most sensitive retinal neurons to diabetes-induced stress, leads to severe visual impairments. Current treatments of DR mainly include intra-vitreal injection of anti-VEGF drugs, laser photocoagulation and vitrectomy. Anti-VEGF agent, currently the mainstay of therapy for both NPDR and PDR, requires persistent injections and only targets retinal neovasculatures. Many DR patients even show inadequate response to anti-VEGF medications after a long period of treatment (5). Thus, fresh insights into DR pathology are needed for identification of novel therapeutic targets.

N^6^-methyladenosine (m^6^A) modification, mainly catalyzed by m^6^A methyltransferase complex (writers), removed by m^6^A demethylases (erasers), and recognized by m^6^A-binding proteins (readers), is one of the most prevalent and abundant internal modifications of mRNA in eukaryotes (6). It regulates mRNA metabolism, including splicing, stability, translation and nuclear export (7–9). Increasing evidence has implied the pathological involvement of aberrant m^6^A modification levels and expression of its modulators (writers, erasers and readers) in DR (10, 11), implying the crucial roles of m^6^A modification in DR pathogenesis. The fat mass and obesity-associated (FTO) protein, which mediates oxidative demethylation of different RNA species, acts as a regulator of fat mass, adipogenesis and energy homeostasis (12–15). Although the clinical association between FTO and diabetes has long been discussed, roles and regulatory networks of FTO in DR remain unclear.

Herein, we revealed that FTO promotes endothelial cell cycle progression and tip cell formation to facilitate angiogenesis in DR. We also found that FTO triggers diabetes-induced microvascular leakage by regulating EC-pericyte crosstalk. FTO also mediated EC-microglia interactions to interrupt retinal homeostasis by inducing microglia activation and neurodegeneration. Mechanistically, FTO regulated diabetic retinal phenotypes through its demethylation activity by modulating *CDK2* mRNA stability with YTH domain-containing family protein 2 (YTHDF2) as the reader. FTO up-regulation in ECs under diabetic conditions was triggered by lactic acid via histone lactylation. FB23-2, which directly binds to FTO and selectively inhibits FTO’s m^6^A demethylase activity, suppressed diabetes induced endothelial phenotypes. We also developed a novel macrophage membrane coated and poly (lactic-co-glycolic acid) (PLGA)-1, 1’-dioctadecyl-3, 3, 3’, 3’-tetramethylindocarbocyanine perchlorate (Dil) based nanoplatform encapsulating FB23-2 for systemic administration. Targeting and therapeutic efficiencies of this nanoplatform have been evaluated and confirmed, indicating its promising role as a treatment agent for DR.

## Results

### Identification of FTO as a potential regulator of DR

M^6^A modification was involved in various pathological processes, while its role in DR is not fully understood. Herein, to determine whether m^6^A levels were altered in DR, we exposed human umbilical vein endothelial cells (HUVECs) to high glucose (25 mM) *in vitro*. For *in vivo* analyses, we used the streptozotocin (STZ)-induced diabetic mice, which develop retinal vascular leakage without neovascularization (16). Both dot blot and m^6^A RNA methylation quantification assays detected reduced m^6^A contents in total RNAs of HUVECs treated with high glucose (**Figures 1A-1B**), as well as in neural retinas collected from STZ mice (**Figures 1C-1D**).

**Figure 1.**
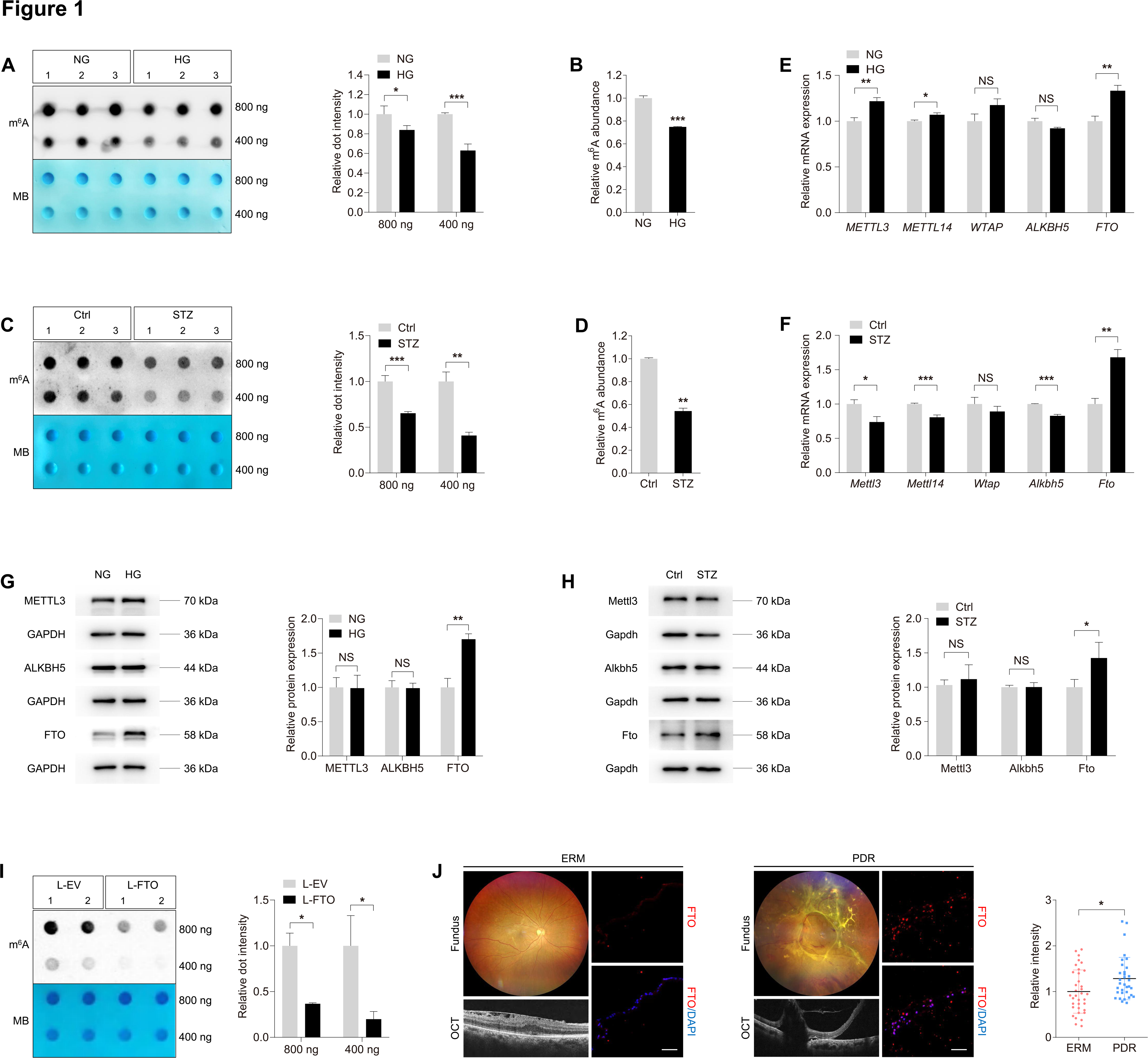
Identification of FTO as a potential regulator of DR. **(A)** m^6^A dot blot assay of global m^6^A abundance in HUVECs treated with normal glucose (5 mM, NG) or high glucose (25 mM, HG) using 800 or 400 ng total RNAs. MB staining is used as a loading control. **(B)** m^6^A RNA methylation quantification assay of global m^6^A abundance in HUVECs treated with NG or HG. **(C)** m^6^A dot blot assay of global m^6^A abundance in retina originated from control or STZ mice using 800 or 400 ng total RNAs. MB staining is applied as a loading control. **(D)** m^6^A RNA methylation quantification assay of global m^6^A abundance in retina originated from control or STZ mice. **(E)** qPCR shows mRNA levels of m^6^A regulators *METTL3*, *METTL14*, *WTAP*, *ALKBH5* and *FTO* in HUVECs treated with NG or HG. **(F)** qPCR presents mRNA levels of m^6^A regulators *Mettl3*, *Mettl14*, *Wtap*, *Alkbh5* and *Fto* in retina originated from control or STZ mice. **(G)** Immunoblotting of METTL3, ALKBH5 and FTO in HUVECs treated with NG or HG. GAPDH is used as an internal control. **(H)** Immunoblotting of Mettl3, Alkbh5 and Fto in retina originated from control or STZ mice. GAPDH is used as an internal control. **(I)** m^6^A dot blot assay of global m^6^A abundance in HUVECs transduced with L-EV or L-FTO using 800 or 400 ng total RNAs. MB staining is applied as a loading control. **(J)** Immunofluorescence staining of FTO in retinal proliferative membranes obtained from DR patients or ERMs isolated from age matched controls without diabetes. Fundus photograph and OCT of DR patients or age matched controls with ERM are shown. NS: not significant (p>0.05); *p<0.05; **p<0.01; and ***p<0.001 (two-tailed Student’s t test). Data are representative of three independent experiments with three biological replicates (mean ± SEM of triplicate assays; **A-I**) or are representative of three independent experiments with similar results (**A**, **C** and **G-I**).

We next aimed to identify the specific m^6^A regulator responsible for the decreased m^6^A levels in high glucose treated HUVECs and STZ retinas. Expression of m^6^A writers (METTL3, METTL14 and WTAP) and erasers (ALKBH5 and FTO) was detected. Both qPCR and immunoblotting analyses demonstrated consistent up-regulation of *FTO* mRNA and protein upon high glucose treatment *in vitro* and *in vivo* (**Figures 1E-1H**). To further test whether FTO directly regulates m^6^A modification in ECs, we overexpressed FTO in HUVECs using lentivirus containing coding sequence (CDS) of human *FTO* gene and tagged with FLAG (L-FTO). Immunoblotting detected FLAG expression and significantly increased FTO expression in HUVECs transduced with L-FTO (**Supplementary Figure S1A**). Immunofluorescence staining further demonstrated that the overexpressed FTO-FLAG fusion protein is mainly located in nuclei (**Supplementary Figures S1B**). M^6^A dot blot assay identified reduced m^6^A level in total RNAs of HUVECs transduced with L-FTO (**Figure 1I**), supporting that FTO suppresses m^6^A modification in HUVECs.

We next aimed to tell the association between FTO and DR using clinical samples. We compared FTO expression in retinal proliferative membranes obtained from DR patients and epi-retinal membranes (ERMs) isolated from age matched controls without diabetes. Immunofluorescence staining revealed stronger FTO intensity in retinal proliferative membranes compared to ERMs (**Figure 1J**), further implying its critical role in DR course.

### FTO promotes endothelial cell cycle progression and tip cell formation to facilitate angiogenesis in vitro

Glucose-mediated microvascular damage is one of the first events in DR (17). Increasing evidence indicates hyperglycemia stimulated retinal vascular EC dysfunction as the pathological basis of DR (18). We therefore initially determined FTO’s roles in regulating EC features *in vitro*. FTO was overexpressed in HUVECs using L-FTO and was knocked down by small interfering RNAs (siRNAs). Three pairs of siRNAs targeting the *FTO* gene (FTO-siRNA) were designed and synthesized, and FTO-siRNA #3 with highest efficiency was selected for further assessments (**Supplementary Figures S1C-S1D**).

RNA transcriptome sequencing (RNA-Seq) was applied to annotate aberrantly changed biological processes and signaling pathways induced by FTO overexpression in HUVECs. A total of 1770 differentially expressed genes [Log_2_ fold change (FC) >0.5 or <0.5; p<0.05], consisting of 771 up-regulated and 999 down-regulated genes, were identified upon FTO overexpression (**Figure 2A**). Both Gene Ontology (GO) and Gene Set Enrichment Analyses (GSEA) revealed that biological processes related to cell cycle progression is enriched upon FTO overexpression (**Figures 2B-2C**), implying the potential role of FTO in endothelial proliferation and angiogenesis. RNA-Seq detected up-regulation of genes regulating all cell cycle phases, including G1, S, G2 and M phases, in HUVECs transduced with L-FTO (**Figures 2D-2E**). Consistently, flow cytometric analyses confirmed that cell cycle, demonstrated by percentage of cells in S and G2/M phases, was accelerated in HUVECs transduced with L-FTO (**Figure 2F**) and was suppressed in cells transfected with FTO-siRNA (**Figure 2G**). 5-Ethynyl-2’-deoxynridine (EdU) assay further confirmed promoted proliferation in HUVECs overexpressing FTO (**Figure 2H**) and inhibited proliferation in cells with FTO knocked down (**Figure 2I**).

**Figure 2.**
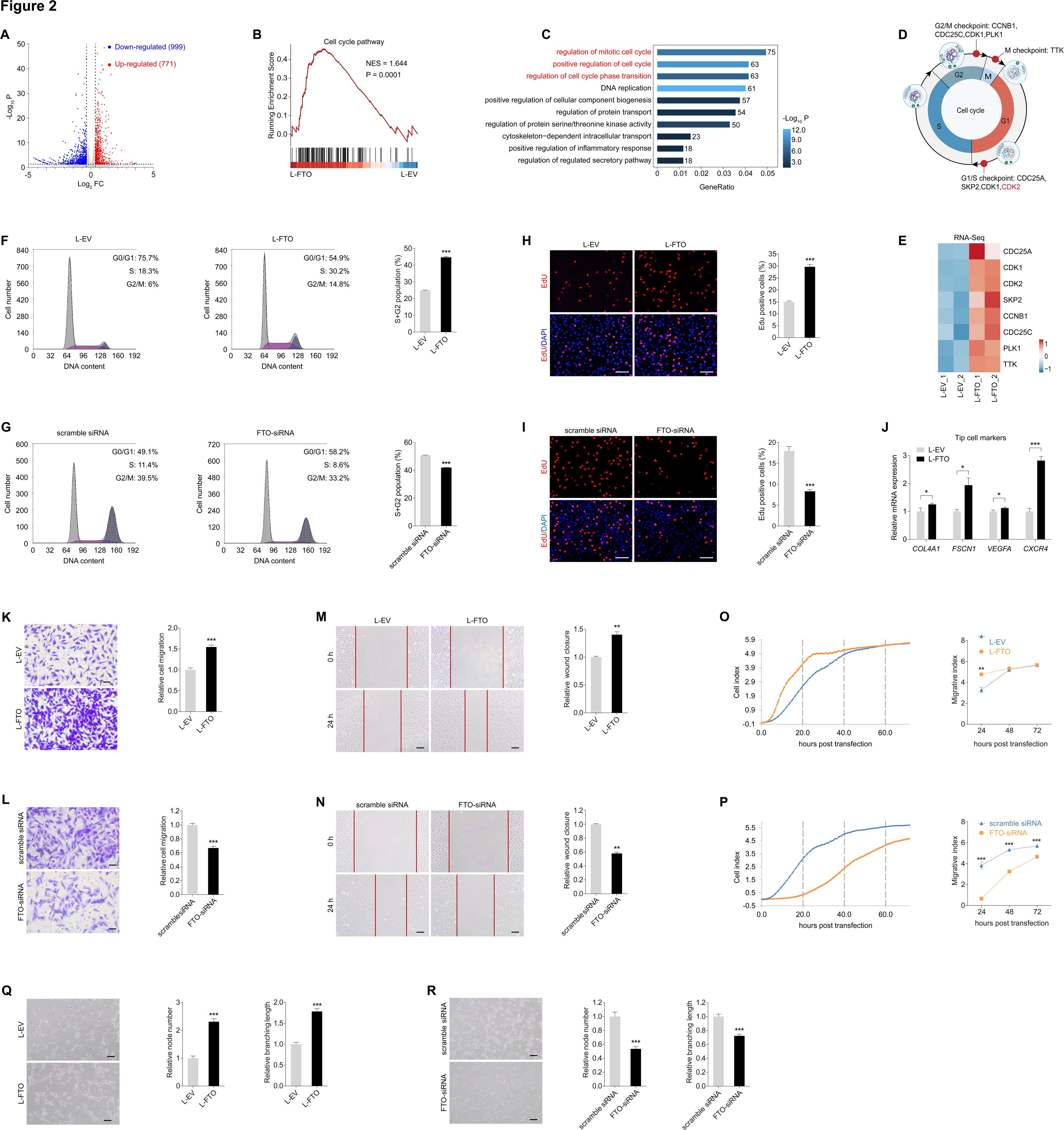
FTO promotes endothelial cell cycle progression and tip cell formation to facilitate angiogenesis *in vitro*. **(A)** Volcano diagram of RNA-Seq data from HUVECs transduced with L-FTO compared to cells transduced with L-EV (Log_2_ FC > 0.5 or < -0.5; *p* < 0.05) is shown. **(B-C)** GSEA and GO plots **(C)** of pathways enriched in HUVECs transduced with L-FTO are presented. **(D-E)** Diagram **(D)** and heatmap **(E)** show differential expression of genes involved in cell cycle checkpoints in HUVECs transduced with L-EV or L-FTO. **(F-G)** Flow cytometric analyses on HUVECs transduced with L-EV/L-FTO **(F)**, or transfected with scramble siRNA/FTO-siRNA **(G)**. **(H-I)** EdU assay on HUVECs transduced with indicated lentivirus **(H)** or transfected with distinct siRNAs **(I)**. Cell nuclei are counterstained with DAPI. Scale bar: 60 µm. **(J)** mRNA levels of tip cell markers (*COL4A1*, *FSCN1*, *VEGFA* and *CXCR4*) detected by qPCR in HUVECs transduced with indicated lentivirus. **(K-L)** Transwell migration assay on HUVECs transduced with indicated lentivirus **(K)** or transfected with distinct siRNAs **(L)**. Scale bar: 50 µm. **(M-N)** Scratch test on HUVECs transduced with indicated lentivirus **(M)** or transfected with distinct siRNAs **(N)**. Scale bar: 100 µm. **(O-P)** RTCA system demonstrates migration rates of HUVECs transduced with L-EV/L-FTO **(O)**, or transfected with scramble siRNA/FTO-siRNA **(P)**. **(Q-R)** Tube formation assay on HUVECs transduced with L-EV/L-FTO **(Q)**, or transfected with scramble siRNA/FTO-siRNA **(R)**. Scale bar: 100 µm. *p<0.05; **p<0.01; and ***p<0.001 (two-tailed Student’s t test). Data are representative of three independent experiments with three biological replicates (mean ± SEM of triplicate assays; **F-R**) or are representative of three independent experiments with similar results (**F-I** and **K-R**).

Endothelial tip cells guide vascularization in neural retina (19). Elevated expression of tip cell markers, including *COL4A1*, *FSCN1*, *VEGFA* and *CXCR4*, was detected in HUVECs overexpressing FTO by qPCR assay, suggesting that FTO promotes endothelial tip cell formation (**Figure 2J**). Effects of FTO on endothelial migration and tube formation were subsequently investigated. As shown by Transwell migration assay and scratch test, cell migration was facilitated in HUVECs transduced with L-FTO (**Figures 2K and 2M**) but was restrained in cells transfected with FTO-siRNA (**Figures 2L and 2N**). Additionally, xCELLigence real-time cell analysis (RTCA) system suggested increased migration rates in HUVECs overexpressing FTO (**Figure 2O**) and decreased rates in cells with FTO knocked down (**Figure 2P**). We also revealed that tube formation, as reflected by node number and branching length, was promoted in HUVECs transduced with L-FTO (**Figure 2Q**) while was inhibited in cells transfected with FTO-siRNA (**Figure 2R**). Collectively, FTO promotes endothelial cell cycle progression and tip cell formation to facilitate angiogenesis *in vitro*.

### FTO promotes endothelial tip cell formation to facilitate neovascularization in mice and zebrafish

We next analyzed whether FTO overexpression associates with neovascularization *in vivo*. FTO protein sequence was highly conserved among human, mice and zebrafish (**Supplementary Figure S2**). Recombinant adeno-associated virus (AAV) containing CDS of mice *Fto* gene with Flag tag and the promoter region of mice Tie2 (AAV-Fto) was constructed and intra-vitreal injected to modulate *Fto* expression in mice vascular ECs. Fto protein level was elevated in neural retina of mice receiving intra-vitreal AAV-Fto injection compared to mice injected with blank AAV containing Flag tag (AAV-blank; **Supplementary Figure S3A**). Expression of Fto-Flag fusion protein was also detected in AAV-FTO injected mice retina (**Supplementary Figure S3B**), and was localized along with the retinal vasculatures (**Supplementary Figure S3C**).

To annotate the role of FTO in regulating mice retinal angiogenesis, we intra-vitreal injected AAV-Fto into mice with oxygen-induced retinopathy (OIR) at P12, the beginning time point of retinal neovascularization and vascular leakage (20). Retina was collected and examined at P17 in OIR, before the regression of pathological vessels (**Figure 3A**). More extensive areas of neovascular tufts (NVTs), formed in the superficial vascular plexuses, were observed in OIR mice receiving AAV-Fto injection (**Figures 3B-3C**). We also noticed that, in the angiogenic area, *Fto* overexpression leads to increased amount of endothelial tip cells (**Figures 3D-3E**). However, no difference in avascular area in the central retina and vessel density in the peripheral deep plexuses was detected between OIR mice injected with AAV-Fto and AAV-blank (**Figures 3B and 3F**). No NVT or avascular area was observed in the control group (**Figure 3B**). Collectively, our data suggested that FTO simultaneously suppressed healthy vascular network formation into the ischemic retina, facilitated pathological NVTs development, and promoted endothelial tip cell formation in the diseased vessel remodeling period of OIR.

**Figure 3.**
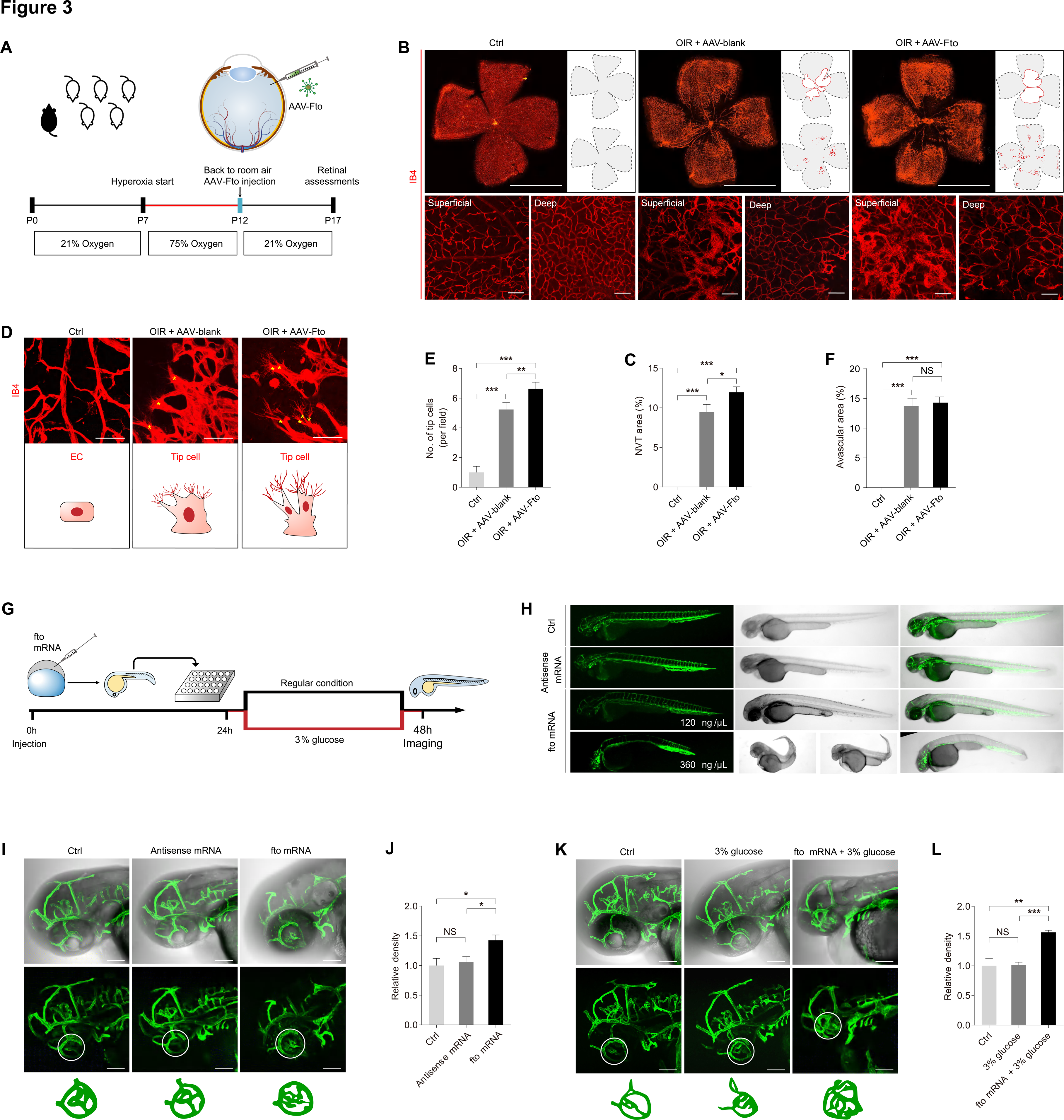
FTO promotes endothelial tip cell formation to facilitate neovascularization in mice and zebrafish. **(A)** Experimental scheme for **(B-F)**. **(B-C)** Fluorescent staining of IB4 in retinal flat mounts originated from control and OIR mice intra-vitreal injected with AAV-blank/AAV-Fto at P17. Magnificent images are shown below to better visualize superficial and deep vascular plexuses. Gray dotted lines indicate the edge of the retina. Red lines suggest the avascular area. NVTs are represented by red dots. Representative images **(B)** along with quantification results of NVT area **(C)** are shown. Scale bar: 2000 µm (up); 50 µm (below). **(D-E)** Fluorescent staining of IB4 in retinal flat mounts of control and OIR mice intra-vitreal injected with AAV-blank/AAV-Fto at P17. Tip cells are indicated by yellow asterisks. Representative images **(D)** along with quantification results of tip cell number per field **(E)** are shown. Scale bar: 50 µm. **(F)** Quantification results of avascular area in retinas collected from mice receiving distinct treatments are shown. **(G)** Experimental scheme for **(H-L)**. **(H)** Truncal vasculatures demonstrated by endogenous EGFP and morphological structures of control and zebrafish injected with antisense or *fto* mRNA at indicated concentrations. **(I-L)** Ocular vasculatures shown by endogenous EGFP and morphological structures of control and zebrafish receiving indicated treatments. Ocular vasculatures are indicated by white circles. Representative images (**I** and **K**) along with quantification results of ocular vessel density (**J** and **L**) are shown. Scale bar: 100 µm. NS: not significant (p > 0.05), *p < 0.05, **p < 0.01, ***p < 0.001 (one-way ANOVA followed by Bonferroni’s test). Data are representative of two to three independent experiments with ten to twelve mice/ four to seven zebrafish per group (mean ± SEM; **C, E-F**, **J** and **L**) or are representative of two or three independent experiments with similar results (**B, D**, **H**, **I** and **K**).

We further analyzed FTO’s effects on mediating vascular functions in zebrafish. We overexpressed FTO in zebrafish through embryonic injection of zebrafish *fto* mRNA in the living transgenic zebrafish strain *Tg(LR57:GFP)*, which contains the enhanced green fluorescent protein (EGFP) cDNA under control of the *fli1* promoter, thus allowing us to visualize its systemic vessels, including ocular vasculatures. Zebrafish embryos injected with *fto* mRNA at the concentration of 120 ng/µL showed no remarkable systemic changes, and were collected and examined at 48 hours post fertilization (**Figures 3G-3H**). Fluorescence staining detected enhanced density of ocular vasculatures in zebrafish injected with *fto* mRNA compared to the uninjected group or those injected with antisense mRNA (**Figure 3I-3J**). To further investigate the role of fto under diabetic condition, fish were cultured in water added with 3% glucose. Consistently, ocular vessel density was increased upon FTO overexpression (**Figures 3K-3L**). These data indicated that FTO promotes zebrafish ocular vascularization under both normal and diabetic conditions. Collectively, we found that FTO regulates endothelial tip cell formation to promote neovascularization in mice and zebrafish.

### FTO regulates EC-pericyte crosstalk and triggers diabetes-induced microvascular dysfunction in mice

To further annotate the role of FTO in regulating diabetes-induced microvascular dysfunction, we assessed retinal vasculatures in STZ mice receiving twice intra-vitreal injections of AAV-Fto (**Figure 4A**). The STZ mice develop retinal vascular leakage without neovascularization (16). Fundus photograph identified cotton wool spot-like lesions in STZ mice injected with AAV-Fto (**Figure 4B**). Fluorescence fundus angiography (FFA), Texas red dextran and Evans blue assays were conducted to further explore FTO’s role on retinal vascular leakage. All three experiments consistently revealed that FTO overexpression aggravated diabetes-induced retinal vascular leakage (**Figures 4C-4E**).

**Figure 4.**
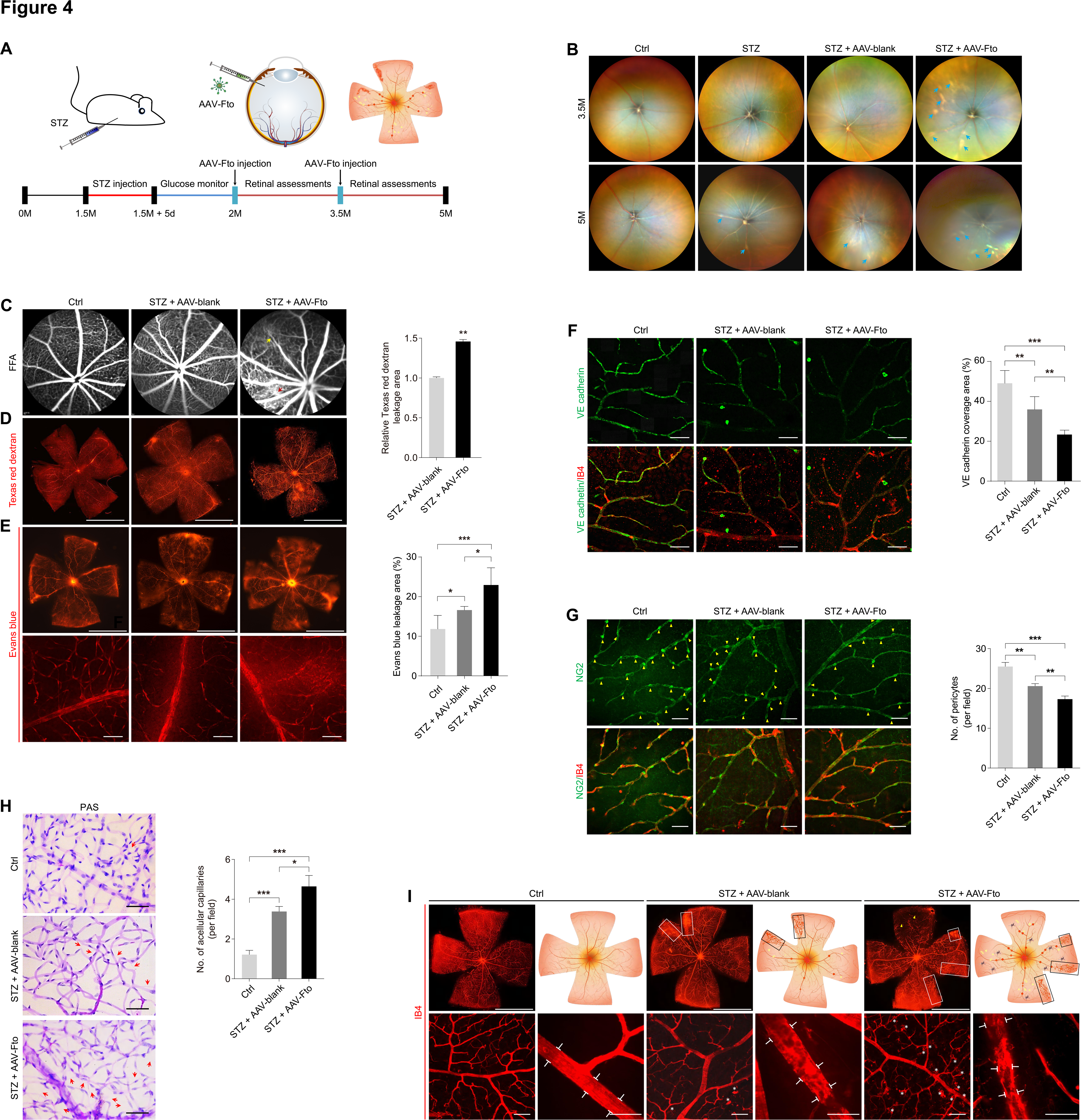
FTO regulates EC-pericyte crosstalk to trigger diabetes-induced microvascular dysfunction in mice. **(A)** Experimental scheme for **(B-I)**. **(B-C)** Representative images of fundus photos **(B)** and FFA of control or STZ mice intra-vitreal injected with AAV-blank/AAV-Fto at indicated time points after injection. Blue arrows represent cotton wool spot-like lesions. Red arrow indicates leakage spot. Yellow arrow suggests abnormal vascular perfusion. **(D-E)** Fluorescence of Texas red dextran **(D)** and Evans blue **(E)** are visualized in retinal flat mounts of control or STZ mice intra-vitreal injected with AAV-blank/AAV-Fto. Scale bar: 2000 µm (**D** and up in **E**); 50 µm (below in **E**). **(F)** Immunofluorescence staining of VE-cadherin and IB4 in retinal flat mounts of control or STZ mice intra-vitreal injected with AAV-blank/AAV-Fto. Scale bar: 50 µm. **(G)** Immunofluorescence staining of NG2 and IB4 in retinal flat mounts of control or STZ mice intra-vitreal injected with AAV-blank/AAV-Fto. Yellow arrowheads indicate pericytes. Scale bar: 50 µm. **(H)** PAS staining of trypsin digested retinal vessels isolated from control or STZ mice intra-vitreal injected with AAV-blank/AAV-Fto. Scale bar: 50 µm. **(I)** Fluorescence staining of IB4 in retinal flat mounts of control or STZ mice intra-vitreal injected with AAV-blank/AAV-Fto. IRMAs are represented by white and black rectangles. Capillary dropout regions are suggested by yellow arrowheads. White brackets indicate structures of main vessels, and white asterisks represent activated microglia cells wrapping around retinal vessels. Scale bar: 2000 µm (up); 50 µm (below). *p < 0.05, **p < 0.01, ***p < 0.001 (two-tailed Student’s t test for **D**; one-way ANOVA followed by Bonferroni’s test for **E-H**). Data are representative of two to three independent experiments with four to eight mice per group (mean ± SEM; **D-H**) or are representative of two or three independent experiments with similar results (**B-I**).

Vascular leakage in the diabetic retina is usually caused by breakdown of the BRB, a biological unit comprised of capillary ECs firmly connected by intercellular tight junctions and their surrounding cells (21). Thus, we stained retinal flat mounts with vascular-endothelial-specific cadherin (VE-cadherin), which maintains the integrity of the EC barrier and attenuates VEGF signaling to suppress angiogenesis (22, 23). VE-cadherin in diabetic retinas was discontinuous compared to non-diabetic animals (**Figure 4F**). Both intensity and coverage area of VE-cadherin were remarkably reduced in STZ mice injected with AAV-Fto compared to those injected with AAV-blank (**Figure 4F**). Pericyte loss in retinal capillaries can induce BRB destruction (24, 25). We further used NG2 and IB4 immunofluorescence staining to detect pericyte coverage of retinal vessels. Consistently, the combination of FTO overexpression and diabetes reduced pericyte coverage (**Figure 4G**).

Vascular lesions, such as acellular capillaries and intraretinal microvascular abnormalities (IRMA), are typical pathological features of diabetic retinas (25). We used trypsin digestion and periodic acid Schiff (PAS) staining to detect FTO associated structural changes in retinal vessels. Increased number of acellular capillaries was detected in STZ mice injected with AAV-Fto compared to both the control group and STZ mice injected with AAV-blank (**Figure 4H**). Additionally, fluorescence staining of IB4 in retinal flat mounts demonstrated enlarged areas of IRMA in STZ mice receiving AAV-Fto injection compared to STZ mice injected with AAV-blank (**Figure 4I**). Uneven and tortuous retinal vasculatures were also observed upon the combination of FTO overexpression and diabetes in mice (**Figure 4I**). Collectively, above data suggested that FTO overexpression and diabetes regulated EC-pericyte crosstalk and aggravated microvascular pathology in a synergistic manner.

### FTO triggers vascular inflammation and regulates EC-microglia crosstalk in vitro

Growing evidence revealed the critical role of retinal inflammation in impaired endothelial function, vascular leakage, pericyte loss and retinal neovascularization (26), we therefore tested whether FTO associates with vascular inflammation. Both RNA-Seq and tandem mass tag (TMT)-based quantitative proteomic analyses detected up-regulation of pro-inflammatory genes/proteins and down-regulation of anti-inflammatory genes/proteins in HUVECs overexpressing FTO (**Figures 5A-5B**). Liquid protein chip (LiquiChip), also known as flexible multi-analyte profiling technology, further identified increased level of pro-inflammatory chemokines (CSF, IL-18 and RANTES) and decreased level of anti-inflammatory cytokines (LIF, IL-4, IL-3 and IL-10) in the culture medium of HUVECs overexpressing FTO (**Figure 5C**). Above data suggested that FTO overexpression in EC triggers vascular inflammation.

**Figure 5.**
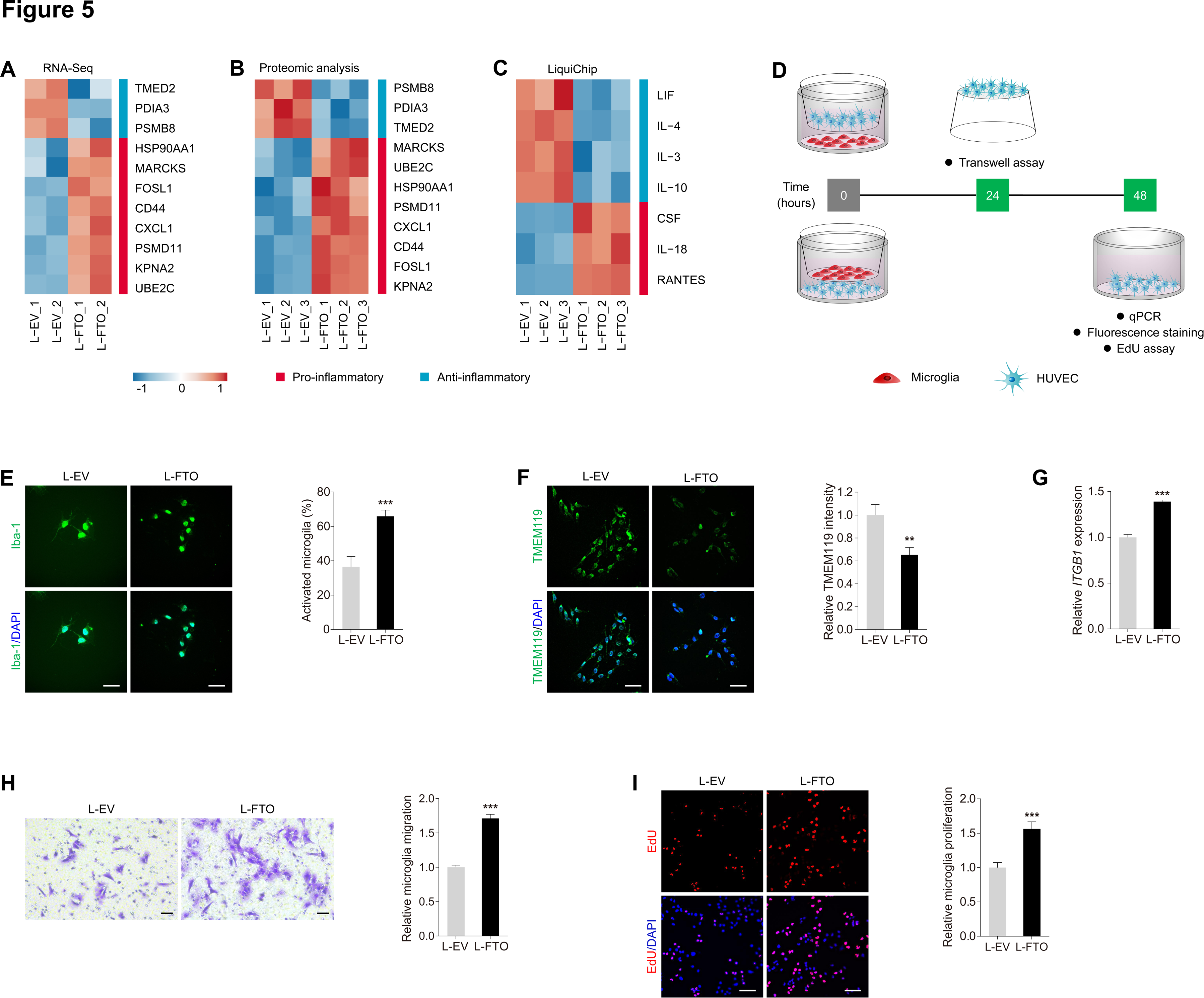
FTO triggers vascular inflammation and regulates EC-microglia crosstalk *in vitro*. **(A-B)** Hierarchical clustering of pro- and anti-inflammatory genes detected by RNA-Seq **(A)** and proteins revealed by TMT-based quantitative proteomic analyses **(B)** in HUVECs transduced with L-EV or L-FTO. **(C)** Heatmap of pro- and anti-inflammatory cytokines in the culture medium of HUVECs transduced with L-EV or L-FTO as detected by LiquiChip. **(D)** Experimental scheme for **(E-I)**. **(E-F)** Immunofluorescence staining of Iba-1 and TMEM119 **(F)** in microglia co-cultured with HUVECs transduced with L-EV or L-FTO. Cell nuclei are counterstained with DAPI. Scale bar: 65 µm. **(G)** mRNA level of *ITGB1* detected by qPCR in microglia co-cultured with HUVECs transduced with L-EV or L-FTO. **(H)** Transwell migration assay on microglia co-cultured with HUVECs transduced with L-EV or L-FTO. Scale bar: 50 µm. **(I)** EdU assay on microglia co-cultured with HUVECs transduced with L-EV or L-FTO. Scale bar: 60 µm. **p < 0.01, ***p < 0.001 (two-tailed Student’s t test). Data are representative of three independent experiments with three biological replicates (mean ± SEM of triplicate assays, **E-I**) or are representative of three independent experiments with similar results (**E-F** and **H-I**).

Vascular inflammation and subsequent microglia activation are typical features of DR and are critical in DR progression (27, 28). To further annotate the role of FTO in regulating EC-microglia crosstalk, we co-cultivated the human microglial clone 3 (HMC3) cells with HUVECs transduced with L-FTO (**Figure 5D**). Immunofluorescence staining of ionized calcium binding adapter molecule 1 (Iba-1), a calcium-binding protein that participates in membrane ruffling and phagocytosis of activated microglia, revealed that percentage of activated microglia, represented by reduced number of interactions of microglia, was increased upon co-cultivation with HUVECs overexpressing FTO (**Figure 5E**). Immunofluorescence staining also identified decreased intensity of transmembrane protein 119 (TMEM119), which specifically labels resident and resting microglia (29, 30), after co-culture with HUVECs transduced with L-FTO (**Figure 5F**).

Activation of microglia cells additionally enhanced their migration and proliferation (17, 27). RNA expression of *ITGB1*, which is responsible for the recruitment and migration of microglia (31), was increased in HMC3 cells co-cultured with HUVECs overexpressing FTO as revealed by qPCR (**Figure 5G**). Transwell migration assay also identified facilitated migration of HMC3 cells sharing medium with HUVECs overexpressing FTO (**Figure 5H**).

Proliferation of HMC3 cells co-cultured with HUVECs transduced with L-FTO was also accelerated as indicated by EdU assay (**Figure 5I**). Collectively, our data suggested that FTO overexpression in EC triggers vascular inflammation and regulates EC-microglia crosstalk to promote microglia activation, migration and proliferation.

### FTO triggers microglia activation and retinal neurodegeneration in mice

Consistent with the above *in vitro* findings, fluorescence staining of IB4 in retinal flat mounts demonstrated accumulation of microglia surrounding the retinal vasculature in STZ mice overexpression FTO (**Figure 4I**). We further asked whether FTO associates with retinal inflammatory response *in vivo*. As revealed by qPCR assay, Fto overexpression aggravated retinal inflammation by elevating expression of *Il1b* and *Ccl2* (**Figure 6A**). We further annotated the role of FTO in regulating microglia features in mice. As revealed by immunofluorescence staining, accumulation of Iba-1 positive microglia was noticed in STZ mice intra-vitreal injected with AAV-Fto (**Figures 6B-6C**).

**Figure 6.**
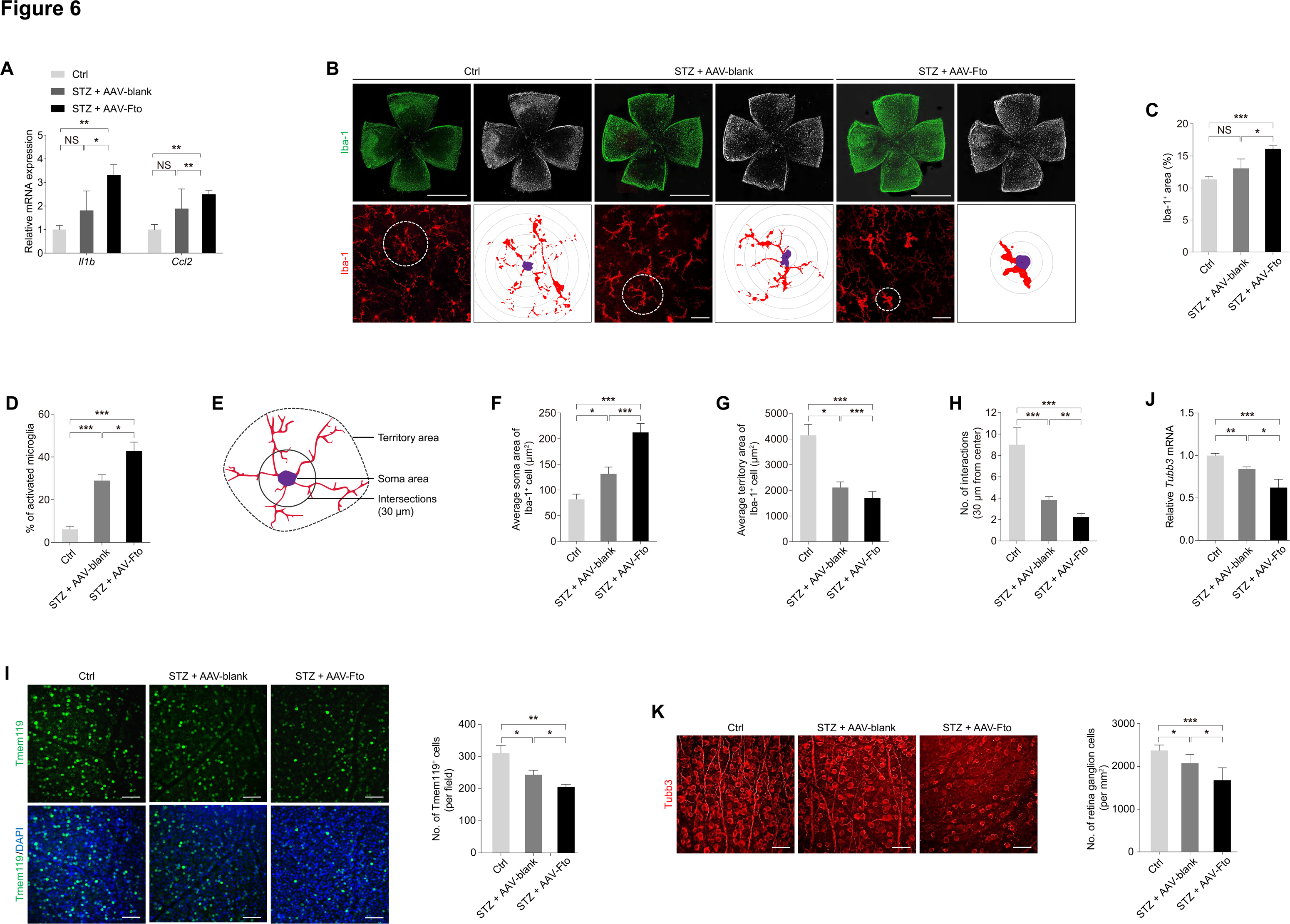
FTO triggers microglia activation and neurodegeneration in mice. **(A)** qPCR shows mRNA levels of inflammatory factors *Il1b* and *Ccl2* in neural retina of control or STZ mice intra-vitreal injected with AAV-blank/AAV-Fto. **(B-D)** Fluorescence staining of Iba-1 in retinal flat mounts of mice receiving indicated treatments is demonstrated. Representative images **(B)** along with quantification of Iba-1 positive microglia area **(C)** and activated microglia **(D)** are shown. Scale bar: 2000 µm (up); 50 µm (below). **(E)** The inset depicts the parameters regarding morphology of microglia. **(F-H)** Quantification results of soma area **(F)**, territory area **(G)** and intersection numbers at 30 µm from nuclei **(H)** in retinal microglia from mice receiving indicated treatments. **(I) F**luorescence staining of Tmem119 in retinal flat mounts of control or STZ mice intra-vitreal injected with AAV-blank/AAV-Fto. Scale bar: 50 µm. **(J)** qPCR demonstrates *Tubb3* mRNA level in neural retina of control or STZ mice intra-vitreal injected with AAV-blank/AAV-Fto. **(K)** Fluorescence staining of Tubb3 in retinal flat mounts of mice receiving indicated treatments is demonstrated. Scale bar: 50 µm. NS: not significant (p > 0.05), *p < 0.05, **p < 0.01, ***p < 0.001 (one-way ANOVA followed by Bonferroni’s test). Data are representative of three independent experiments with three to six mice per group (mean ± SEM; **A**, **C-D** and **F-K**) or are representative of two or three independent experiments with similar results (**B**, **I** and **K**).

Further morphological assessments identified that endothelial FTO overexpression and diabetes promote microglia activation, represented by an amoeboid morphology, in a synergistic manner (**Figures 6B and 6D**). Endothelial FTO overexpression and diabetes also synergistically increased the soma area as well as decreased the territory projection area and number of interactions of microglia (**Figures 6B and 6E-6H**). These morphological changes are consistent with a decreased Tmem119 intensity in neural retina of STZ mice intra-vitreal injected with AAV-Fto, indicating that endothelial FTO overexpression induces microglia activation (**Figure 6I**).

Activation of microglia severely affected retinal neurons, leading to neurodegeneration and vision loss (23). We thus tested whether FTO overexpression was accompanied with RGC loss using the RGC specific marker tubulin beta-III (TUBB3) (32). We found that diabetes and FTO overexpression decreased mRNA level of *Tubb3* in mice neural retina in a synergistic manner (**Figure 6J**). Axonal degeneration and reduced number of Tubb3 positive RGCs were also noticed in neural retinas of STZ mice injected with AAV-Fto (**Figure 6K**). Thus, our data indicated that FTO associates with retinal inflammation, microglia activation and neurodegeneration in mice.

### Demethylation activity is required for FTO to regulate EC features

To annotate whether effects of FTO on EC function depend on its demethylation activity, we introduced two catalytically inactive mutations, H231A and D233A (12, 33), into L-FTO to generate L-FTO^MU^. Dot blot assay indicated that the two mutations remarkably inhibited demethylation activity of FTO (**Figure 7A**), but immunoblotting suggested that FTO protein expression was not affected (**Figure 7B**). Flow cytometric analyses demonstrated accelerated cell cycle process in HUVECs overexpressing wild type FTO protein, but was not affected in cells overexpressing mutant FTO compared to the control group (**Figure 7C**), supporting that the mutations reverse the promotive role of FTO on endothelial cell cycle progression. Facilitated proliferation, indicated by EdU positive cells, was identified in HUVECs transduced with L-FTO^WT^, but was not found in cells infected with L-FTO^MU^ (**Figure 7D**). Moreover, cell migration, demonstrated by Transwell migration assay and scratch test, was facilitated in HUVECs transduced with L-FTO^WT^ but not the mutant form (**Figures 7E-7F**), suggesting that mutant FTO protein does not associate with EC migration.

**Figure 7.**
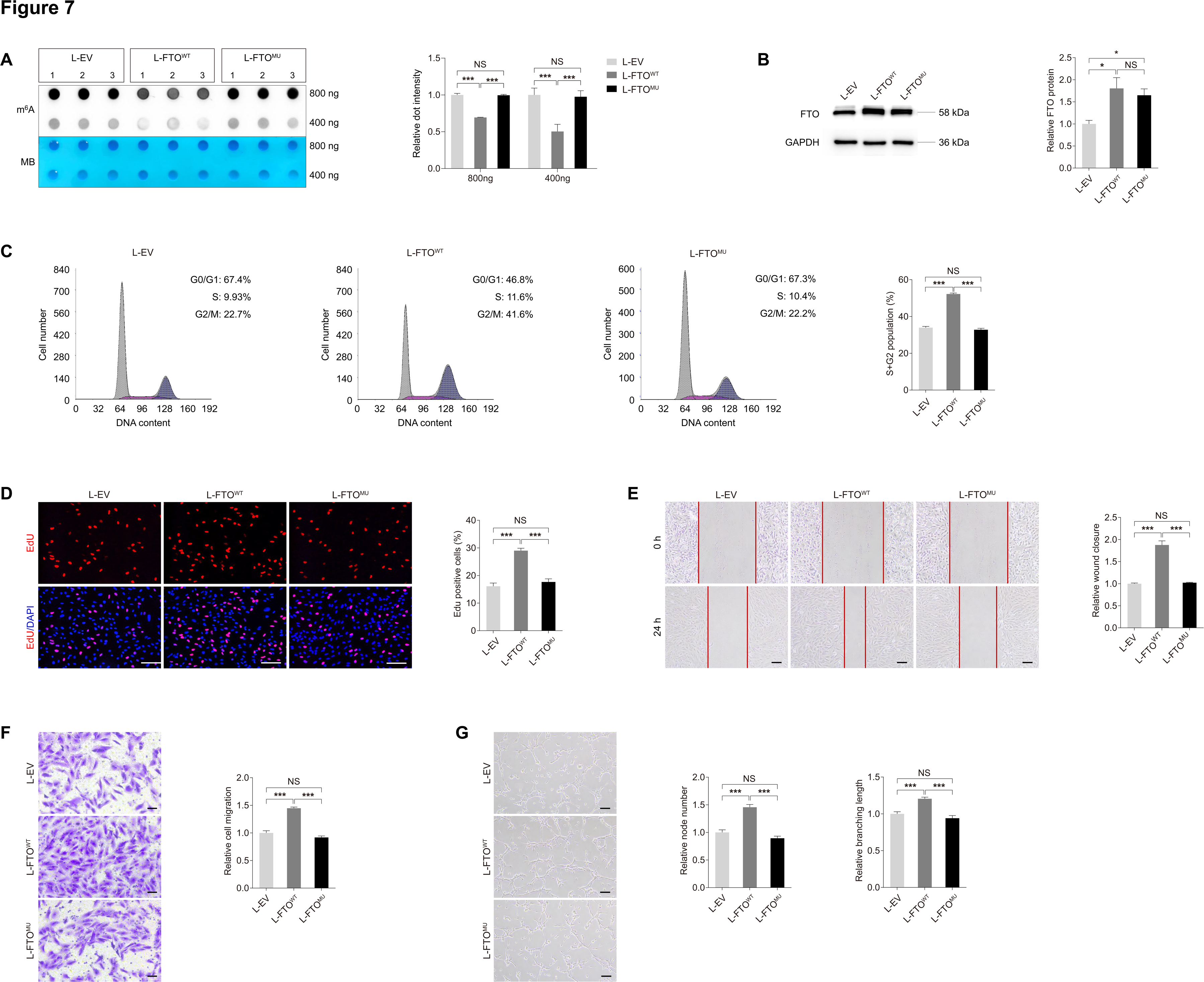
Demethylation activity is required for FTO to regulate EC features. **(A)** m^6^A dot blot assay of global m^6^A abundance in HUVECs transduced with L-EV, L-FTO^WT^ or L-FTO^MU^ using 800 or 400 ng total RNAs. MB staining is used as a loading control. **(B)** Immunoblotting of FTO in HUVECs transduced with L-EV, L-FTO^WT^ or L-FTO^MU^. GAPDH is used as an internal control. **(C)** Flow cytometric analyses of cell cycle process in HUVECs transduced with L-EV, L-FTO^WT^ or L-FTO^MU^. **(D)** EdU assay on HUVECs transduced with indicated lentivirus. Scale bar: 60 µm. **(E)** Scratch test on HUVECs transduced with L-EV, L-FTO^WT^ or L-FTO^MU^. Scale bar: 100 µm. **(F)** Transwell migration assay on HUVECs transduced with indicated lentivirus. Scale bar: 50 µm. **(G)** Tube formation analyses on HUVECs transduced with L-EV, L-FTO^WT^ or L-FTO^MU^. Scale bar: 100 µm. NS: not significant (p>0.05); *p<0.05 and ***p<0.001 (one-way ANOVA followed by Bonferroni’s test). Data are representative of three independent experiments with three biological replicates (mean ± SEM of triplicate assays; **A-G**) or are representative of three independent experiments with similar results (**A-G**).

Tube formation assay also revealed increased node number and branching length in HUVECs transduced with L-FTO^WT^ but not L-FTO^MU^ (**Figure 7G**). Collectively, the effects of FTO on EC will be abolished by introduction of the H231A and D233A mutations. Intact demethylation activity of FTO was required for its regulations on EC features and its pathogenic roles in microvascular dysfunctions.

### FTO regulates CDK2 mRNA stability with YTHDF2 as the reader in an m6A-dependent manner

Since FTO mediates EC features through its demethylation activity, we then applied methylated RNA immunoprecipitation-sequencing (MeRIP-Seq) to identify down-stream targets of FTO in HUVECs. A total of 465 m^6^A-hypo peaks and 586 m^6^A-hyper peaks (Log_2_ FC >0.2 or <-0.2; *p* < 0.05) were initially identified in HUVECs overexpressing FTO compared to cells transduced with L-EV (**Figure 8A**). MeRIP-Seq revealed a dominant distribution of m^6^A peaks in mRNAs (**Figure 8B**), especially in CDS and 3’-untranslated region (3’-UTR) of RNA transcripts (**Figure 8C**). Consistent with previous reports (34, 35), the m^6^A sites displayed the RRACH motif (**Figure 8D**). Given the demethylation activity of FTO, we focused on genes with m^6^A-hypo peaks in HUVECs upon FTO overexpression. We analyzed whether protein expression of these genes was altered using the proteomic data. A total of 9 genes containing m^6^A-hypo peaks with altered protein expression were sorted out (**Figure 8E**). Among all, the *CDK2* gene, encoding a serine/threonine protein kinase that participates in cell cycle regulation, was found involved in cell proliferation and DR (25, 36). MeRIP-Seq data showed that FTO overexpression suppresses the m^6^A level of a peak containing an m^6^A site (Chr1: 56365520-56365521) in the *CDK2* transcript in HUVECs (**Figure 8F**). We also detected elevated *CDK2* mRNA and protein expression in HUVECs exposed to high glucose (**Figures 8G-8H**) and neural retinas of STZ mice (**Figures 8I-8J**). Thus, we emphasized on CDK2 as a potential key regulator of FTO in subsequent studies.

**Figure 8.**
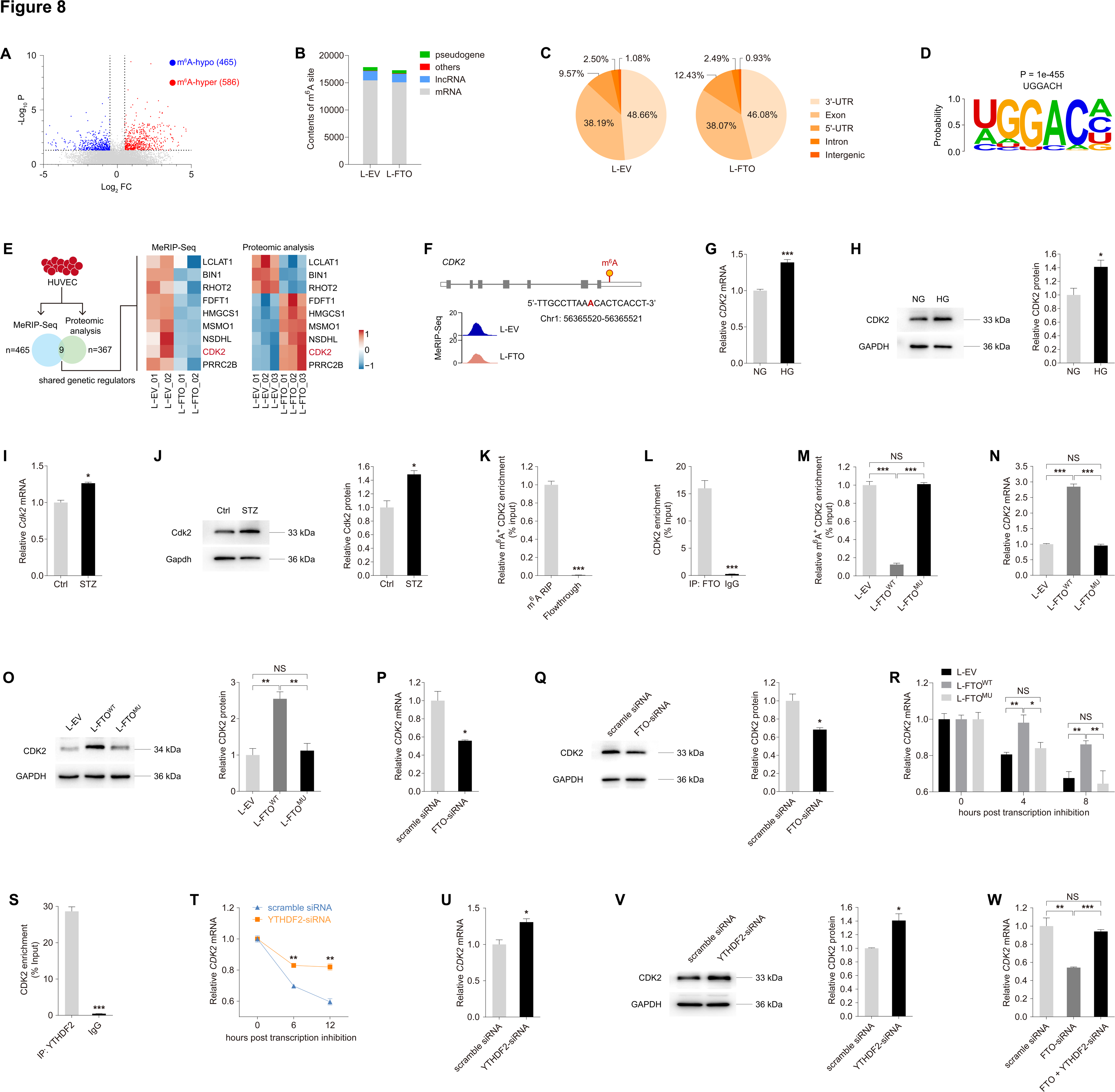
FTO regulates CDK2 mRNA stability with YTHDF2 as the reader in an m^6^A-dependent manner. **(A-B)** Volcano diagram of m^6^A-hypo and m^6^A-hyper peaks identified in HUVECs overexpressing FTO compared to cells transduced with L-EV (Log_2_ FC > 0.2 or < -0.2; *p* < 0.05). **(B-C)** MeRIP-Seq data of HUVECs transduced with L-EV or L-FTO. Contents of m^6^A site in all types of RNAs **(B)** and distribution of total m^6^A peaks in distinct regions of mRNA transcripts **(C)** are presented. **(D)** Sequence of enriched motif displayed by HOMER. **(E)** Genes containing m^6^A-hypo peaks with altered protein expression are sorted out. **(F)** Abundance of m^6^A peak in the *CDK2* transcript in HUVECs transduced with L-EV or L-FTO detected by MeRIP-Seq. Genomic location of its containing m^6^A site is annotated. **(G-H)** mRNA **(G)** and protein **(H)** expression of CDK2 in HUVECs exposed to NG or HG. GAPDH is used as an internal control. **(I-J)** Cdk2 mRNA and protein levels detected in neural retinas of control and STZ mice. Gapdh is used as an internal control. **(K)** MeRIP-qPCR analysis of m^6^A enrichment on *CDK2* mRNA in HUVECs. **(L)** FTO-RIP-qPCR validates the binding between FTO protein and *CDK2* mRNA in HUVECs. **(M)** MeRIP-qPCR analysis of m^6^A enrichment on *CDK2* mRNA in HUVECs transduced with L-EV, L-FTO^WT^ or L-FTO^MU^. **(N-O)** mRNA **(N)** and protein **(O)** expression of CDK2 in HUVECs transduced with indicated lentivirus. GAPDH is used as an internal control. **(P-Q)** CDK2 mRNA **(P)** and protein **(Q)** levels in HUVECs transfected with scramble siRNA or FTO-siRNA. GAPDH was used as an internal control. **(R)** *CDK2* mRNA levels detected by qPCR in HUVECs transduced with indicated lentivirus at 0, 4 and 8 hours post actinomycin D treatment. **(S)** YTHDF2-RIP-qPCR validation of YTHDF2 binding to *CDK2* mRNA in HUVECs. **(T)** *CDK2* mRNA levels detected by qPCR in HUVECs transfected with scramble siRNA or YTHDF2-siRNA at 0, 6 and 12 hours post actinomycin D treatment. **(U-V)** CDK2 mRNA **(U)** and protein **(V)** levels in HUVECs transfected with scramble siRNA or YTHDF2-siRNA. GAPDH is used as an internal control. **(W)** *CDK2* mRNA levels detected by qPCR in HUVECs transfected with scramble siRNA, FTO-siRNA, or FTO- and YTHDF2-siRNA. NS: not significant (p>0.05); *p<0.05; **p<0.01; and ***p<0.001 (two-tailed Student’s t test for **G-L**, **P-Q** and **S-V**, one-way ANOVA followed by Bonferroni’s test for **M-O**, **R** and **W**). Data are representative of three independent experiments with three biological replicates (mean ± SEM of triplicate assays, **G-W**), or are representative of three independent experiments with similar results (**H**, **J**, **O**, **Q** and **V**).

We next investigated whether CDK2 was a direct target of FTO in HUVECs. MeRIP-qPCR confirmed the direct binding between m^6^A antibody and the m^6^A site within the *CDK2* transcript in HUVECs (**Figure 8K**). RNA immunoprecipitation (RIP) assay further validated the binding between the FTO protein and the *CDK2* mRNA (**Figure 8L**). In line with the MeRIP-Seq data (**Figures 8E-8F**), MeRIP-qPCR identified suppressed m^6^A level in the *CDK2* transcript in HUVECs overexpressing wild type FTO, but not its mutant form (**Figure 8M**), suggesting the inhibitory role of FTO on m^6^A contents in CDK2. As revealed by qPCR and immunoblotting, both mRNA and protein expression of CDK2 were increased in HUVECs upon transduction of L-FTO^WT^, while such change was abolished in cells transduced with L-FTO^MU^ (**Figures 8N-8O**). Consistently, CDK2 mRNA and protein levels were decreased in HUVECs transfected with FTO-siRNA (**Figures 8P-8Q**). Collectively, our data implied that FTO regulates CDK2 expression through its demethylation on the *CDK2* transcript.

We further annotated the specific m^6^A reader that binds to CDK2 in EC. Above data suggested that FTO decreased m^6^A level but increased mRNA expression of CDK2, therefore YTHDF2, a well-recognized m^6^A reader that promotes targeted mRNA decay (8, 37), is identified as a potential binding protein to CDK2 in EC. We next investigated whether FTO promotes CDK2 expression by enhancing its mRNA stability with YTHDF2 as a reader. Actinomycin D was used to inhibit CDK2 transcription in HUVECs. Prolonged half-life of the *CDK2* transcript was detected in cells transduced with L-FTO^WT^, but not in cells overexpressing the mutant FTO protein (**Figure 8R**), supporting that FTO intensifies *CDK2* mRNA stability via its demethylation activity. RIP assay further confirmed the direct binding between the YTHDF2 protein and the *CDK2* transcript (**Figure 8S**). We then analyzed whether YTHDF2 knocking down alleviates its mediated decay of the *CDK2* mRNA. Among all three pairs of siRNAs targeting the *YTHDF2* gene, YTHDF2-siRNA #2 presented highest efficiency and was chosen for the following studies (**Supplementary Figures 1E-1F**). Stability of the *CDK2* mRNA was enhanced upon YTHDF2 knocking down (**Figure 8T**), followed by increased expression of the *CDK2* mRNA and protein (**Figures 8U-8V**). We also found that YTHDF2 knocking down could restore the reduced *CDK2* mRNA level induced by FTO-siRNA in HUVECs (**Figure 8W**). Collectively, these findings indicated that FTO regulates *CDK2* mRNA stability in an m^6^A-YTHDF2-dependent manner.

### Lactic acid regulates FTO expression via histone lactylation

We continued to analyze the potential upstream mechanisms responsible for the enhanced FTO expression in DR. Disturbed lactate homeostasis in retina is a common feature of DR (38), we thus aimed to ascertain whether lactate is responsible for the diabetes-driven FTO up-regulation. Increased lactate concentration was detected in HUVECs exposed to high glucose (**Figure 9A**). Addition of lactate into the culture medium of HUVECs up-regulated both mRNA and protein expression of FTO and CDK2 (**Figures 9B-9C**), implying that lactate is potentially responsible for FTO up-regulation in HUVEC.

**Figure 9.**
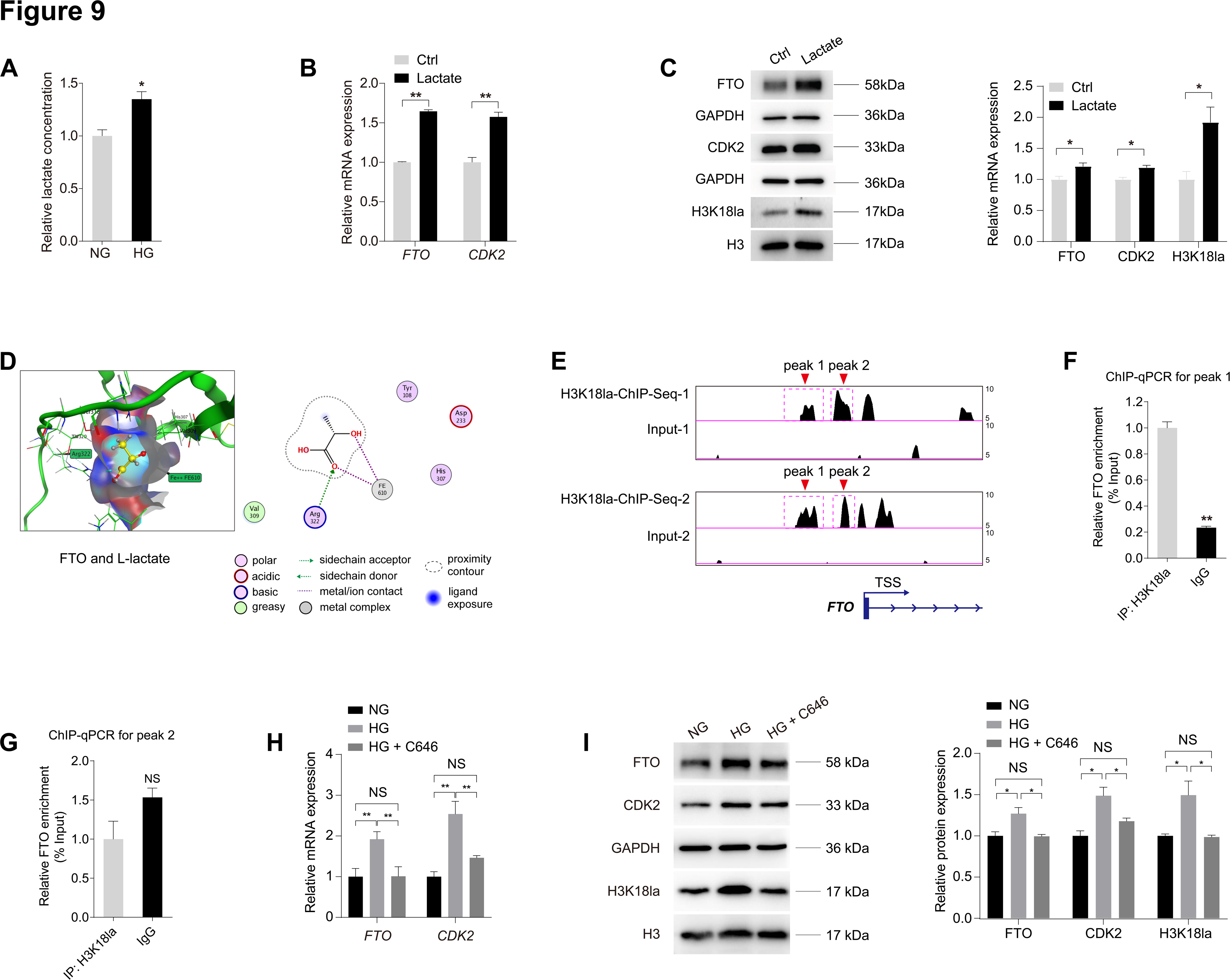
Lactic acid regulates FTO expression via histone lactylation. **(A)** Relative lactate concentration in HUVECs treated with NG or HG. **(B)** qPCR presents mRNA levels of *FTO* and *CDK2* in HUVECs treated with or without lactate (10 mM). **(C)** Immunoblotting of FTO, CDK2 and H3K18la in HUVECs treated with or without lactate. GAPDH and H3 are used as internal controls. **(D)** MOE is used to predict the binding affinity between the FTO protein and L-lactate. **(E)** Enrichment of the H3K18la signal (two peaks) in the promoter region of the *FTO* gene is demonstrated by ChIP-Seq using anti-H3K18la antibodies (GEO accession number: GSE156675). **(F-G)** ChIP-qPCR using anti-H3K18la antibodies validates H3K18la enrichment in peak 1 but not peak 2 within the promoter region of FTO in HUVECs. **(H)** qPCR presents mRNA levels of *FTO* and *CDK2* in HUVECs treated with NG, HG, as well as HG and C646 (100 µM). **(I)** Immunoblotting of FTO, CDK2 and H3k18la in HUVECs with different treatments. GAPDH and H3 are used as internal controls. NS: not significant (p>0.05); *p<0.05; **p<0.01 (two-tailed Student’s t test in **A-C** and **F-G**, One-way ANOVA in **H-I**). Data are representative of three independent experiments with three biological replicates (mean ± SEM of triplicate assays; **A-C** and **F-I**) or are representative of three independent experiments with similar results (**C** and **I**).

We further explored the modification pattern of lactate in HUVECs. We asked whether lactate mediates FTO expression through direct lactylation of FTO or histone lactylation. We used molecular docking in the molecular operating environment (MOE) to predict the binding affinity between L-lactate molecules and the FTO protein. However, no lysine site in the FTO protein was predicted to bind with lactate (**Figure 9D**). Next, we tried to verify whether lactate-H3K18la pathway facilitates FTO expression. Elevated H3K18la level was detected in HUVECs added with lactate (**Figure 9C**). Chromatin immunoprecipitation followed by sequencing (ChIP-Seq) using anti-H3K18la antibodies (GEO accession number: GSE156675) demonstrated a significant enrichment of the H3K18la signal (two peaks) in the promoter region of the *FTO* gene (**Figure 9E**). ChIP-qPCR assay further validated that H3K18la was enriched in peak 1 (**Figures 9F-9G**). Acetyltransferase p300 is a histone Kla “writer” enzyme (39). Notably, we found the elevated FTO, CDK2 and H3K18la levels in HUVECs treated with high glucose were reduced after addition of the p300 inhibitor C646 (**Figures 9H-9I**). Collectively, the above data implied that lactic acid regulates FTO expression via H3K18la.

### FB23-2 suppresses demethylation activity of FTO to inhibit diabetes induced endothelial phenotypes in vitro

FB23-2, which directly binds to FTO and selectively inhibits FTO’s m^6^A demethylase activity, is reported to exhibit FTO-dependent anti-proliferation activity via up-regulating global m^6^A levels (40). We thus tested the effect of FB23-2 on FTO inhibition in HUVECs using dot blot assay. We found that FB23-2 suppresses the demethylation activity of FTO in HUVECs without affecting its protein expression (**Figures 10A-10B**). We next detected whether FB23-2 could rescue the high glucose induced pathogenic endothelial phenotypes. EdU assay implied that the high glucose associated proliferation of HUVECs was partly restrained by FB23-2 treatment (**Figure 10C**). FB23-2 also alleviated the accelerated HUVEC migration caused by high glucose (**Figure 10D**). Moreover, high glucose induced tube formation of HUVECs, as revealed by increased node number and branching length, was restored by the addition of FB23-2 (**Figure 10E**). Collectively, our data suggested that the FTO inhibitor, FB23-2, is a potential agent for diabetes correlated pathogenic endothelial phenotypes.

**Figure 10.**
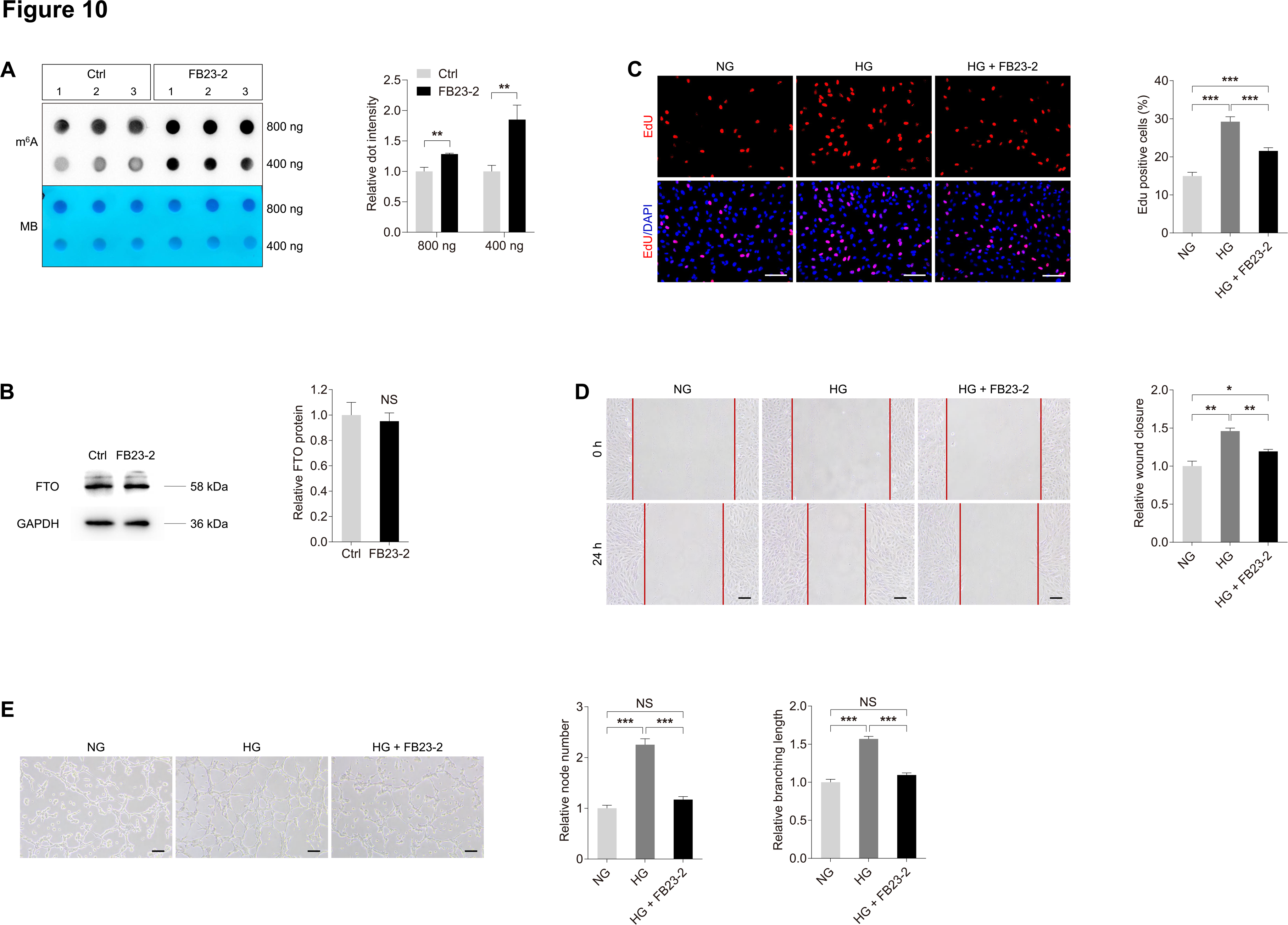
FB23-2 suppresses demethylation activity of FTO to inhibit diabetes induced endothelial phenotypes *in vitro.* **(A)** m^6^A dot blot assay of global m^6^A abundance in HUVECs treated with or without FB23-2 (2 µM) using 800 or 400 ng total RNAs. MB staining is used as a loading control. **(B)** Immunoblotting of FTO in HUVECs treated with or without FB23-2. GAPDH is used as an internal control. **(C)** EdU assay on HUVECs treated with NG, HG, as well as HG and FB23-2. Scale bar: 60 µm. **(D)** Scratch test on HUVECs receiving indicated treatments. Scale bar: 100 µm. **(E)** Tube formation analyses on HUVECs treated with NG, HG, as well as HG and FB23-2. Scale bar: 100 µm. NS: not significant (p>0.05); *p<0.05, **p<0.01 and ***p<0.001 (two-tailed Student’s t test in A-B, One-way ANOVA in **C-E**). Data are representative of three independent experiments with three biological replicates (mean ± SEM of triplicate assays; **A-E**) or are representative of three independent experiments with similar results (**A-E**).

### Characteristics of NP-FB23-2 and evaluation of its therapeutic efficacy on retinal neovascularization in mice

To further explore the therapeutic potential of FB23-2 in DR, we developed a novel macrophage membrane coated and PLGA-Dil based nanoplatform encapsulating FB23-2 for systemic administration (**Figure 11A**). Considering the poor solubility of FB23-2 in water, the polymer cores were prepared using PLGA-Dil as a hydrophobic and fluorescent drug carrier to encapsulate FB23-2. Macrophage membrane coated nanoparticles exhibited significantly enhanced accumulation in retinal neovascular lesions (41), we thereby wrapped the PLGA-Dil-FB23-2 particle in macrophage membrane-derived vesicles (M-vesicles) to get the M-FB23-2 (**Figure 11A**). In addition, the RGD peptide, a vasculature-targeting tri-peptide motif containing arginine, glycine and aspartic acid, was further linked to M-FB23-2 using 1,2-Distearoyl-sn-glycero-3-phosphoethanolamine (DSPE) and polyethylene glycol (PEG) to obtain NP-FB23-2 (**Figure 11A**). Transition electron microscopy (TEM) demonstrated that NP-FB23-2 was spherical in shape with a core-shell structure, indicating the successful cloaking of PLGA-Dil-FB23-2 with M-vesicles (**Figure 11B**). Dynamic light scattering (DLS) measurements demonstrated that NP-FB23-2s show narrow size distributions with the average diameter of 201.71 ± 22.34 nm (**Figure 11C**).

**Figure 11.**
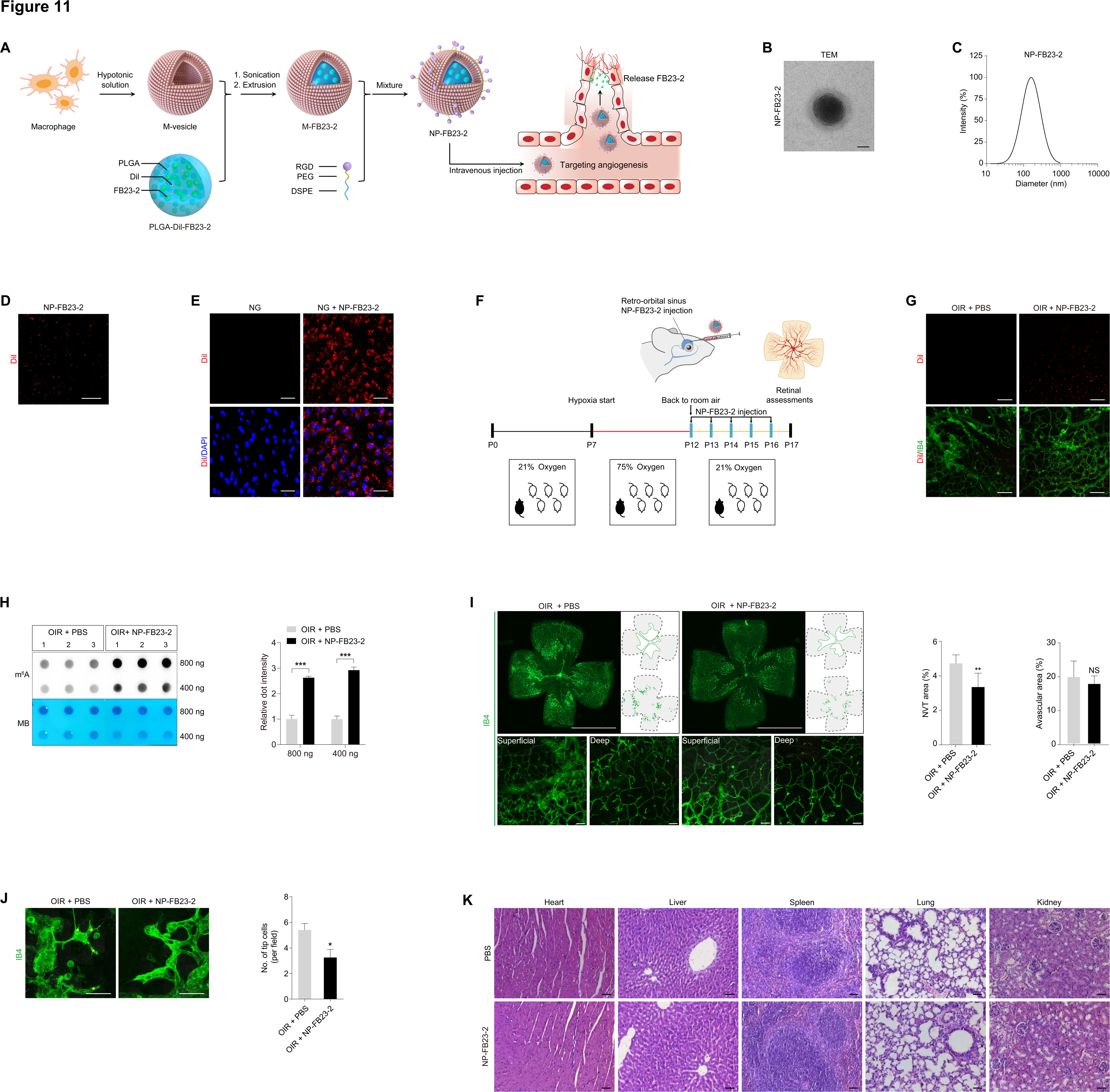
Characteristics of NP-FB23-2 and evaluation of its therapeutic efficacy on retinal neovascularization in mice. **(A)** Experimental scheme illustrating the synthetic procedures of NP-FB23-2. **(B)** TEM of Dil labeled NP-FB23-2. Scale bar: 50 nm. **(C)** The size distribution profile of NP-FB23-2s. Fluorescence demonstrated by Dil labeled NP-FB23-2s. Scale bar: 20 µm. Fluorescent images of HUVECs treated with PBS or NP-FB23-2s. Cell nuclei are counterstained with DAPI. Scale bar: 65 µm. **(F)** Experimental scheme for **(G-J)**. **(G)** Fluorescent images of neural retinas collected from mice intravenously injected with PBS or NP-FB23-2s. Retinal vasculatures are stained with IB4. Scale bar: 50 µm. **(H)** m^6^A dot blot assay of global m^6^A abundance in retinas of OIR mice intravenously injected with PBS or NP-FB23-2s at P17 using 800 or 400ng total RNAs. MB staining is applied as a loading control. **(I)** Fluorescence staining of IB4 in retinal flat mounts originated from OIR mice intravenously injected with PBS or NP-FB23-2s at P17. Superficial and deep vascular plexuses are shown by magnificent images. Gray dotted lines indicate the edge of the retina. Red lines suggest the avascular area. NVTs are represented by red dots. Representative images along with quantification results of NVT and avascular areas are shown. Scale bar: 2000 µm (up); 50 µm (below). **(J)** Fluorescence staining of IB4 in retinal flat mounts originated from OIR mice with indicated treatments at P17. Tip cells are represented by yellow asterisks. Scale bar: 50 µm. **(K)** Representative H&E staining images of major organs including the heart, liver, spleen, lung and kidney from mice intravenously injected with PBS or NP-FB23-2s. *p < 0.05 and ***p < 0.001 (two-tailed Student’s t test). Data are representative of two to three independent experiments with three to seven mice per group (mean ± SEM; **H-J**) or are representative of two or three independent experiments with similar results (**B**, **D-E** and **G-K**).

The cellular uptake of NP-FB23-2s into HUVECs was then examined. As revealed by fluorescence microscopy, the Dil labeled NP-FB23-2s presented red fluorescence (**Figure 11D**). Fluorescence microscopy demonstrated that NP-FB23-2s exhibited strong fluorescent signal inside the HUVECs (**Figure 11E**). Targeting properties of NP-FB23-2s were further evaluated in OIR mice intra-retro-orbital sinus injected with NP-FB23-2s (**Figure 11F**). Fluorescent signal of NP-FB23-2s was enriched in retinal vessels of OIR mice (**Figure 11G**). No fluorescent signal was detected in OIR mice intravenously injected with phosphate buffer saline (PBS; **Figure 11G**). Our data demonstrated the *in vivo* retinal neovasculature-targeting capacity of NP-FB23-2.

We next verified the therapeutic efficacy of NP-FB23-2s in OIR mice intravenously injected with NP-FB23-2s. M^6^A dot blot assay identified increased m^6^A level in total RNAs of OIR neural retinas from mice injected with NP-FB23-2s compared to PBS (**Figure 11H**), implying that NP-FB23-2 alleviates the m^6^A modification reduction in OIR retinas. We also noticed that OIR induced NVTs, formed in the superficial vascular plexuses, are suppressed upon NP-FB23-2 injection (**Figure 11I**). OIR associated increased amount of endothelial tip cells in the angiogenic area was reduced by intravenous NP-FB23-2 injection (**Figure 11J**). However, no difference in avascular area in the central retina and vessel density in the peripheral deep plexuses was detected between OIR mice injected with PBS and NP-FB23-2 (**Figure 11I**). Systemic toxicity of intravenous NP-FB23-2 administration was further evaluated using Hematoxylin and Eosin (H&E) staining. No significant toxicity to major organs including heart, liver, spleen, lung and kidney was exhibited in mice intravenously injected with NP-FB23-2 (**Figure 11K**). Collectively, the above data confirmed the therapeutic efficacy of intravenous NP-FB23-2 injection on retinal neovascularization in mice.

## Discussion

Accumulating evidence suggests that dysregulation of m^6^A modulators participates in the occurrence and progression of DR (10, 42–44), while the roles and regulatory network of FTO in DR have never been elucidated. Herein, we detected elevated expression of RNA demethylase FTO in DR. Both *in vivo* and *in vitro* studies revealed that FTO overexpression in vascular ECs contributes to DR phenotypes, including angiogenesis, vascular leakage, inflammation and neurodegeneration (**Figure 12**). Further assessments validated *CDK2* as a contributor to the FTO induced retinal phenotypes, and lactate-H3K18la pathway as the up-stream regulator of FTO under diabetic conditions. We also developed a novel macrophage membrane coated and PLGA-Dil based nanoplatform encapsulating FB23-2, an FTO inhibitor that suppresses diabetes associated endothelial phenotypes, thus providing a promising nanotherapeutic approach for DR.

**Figure 12.**
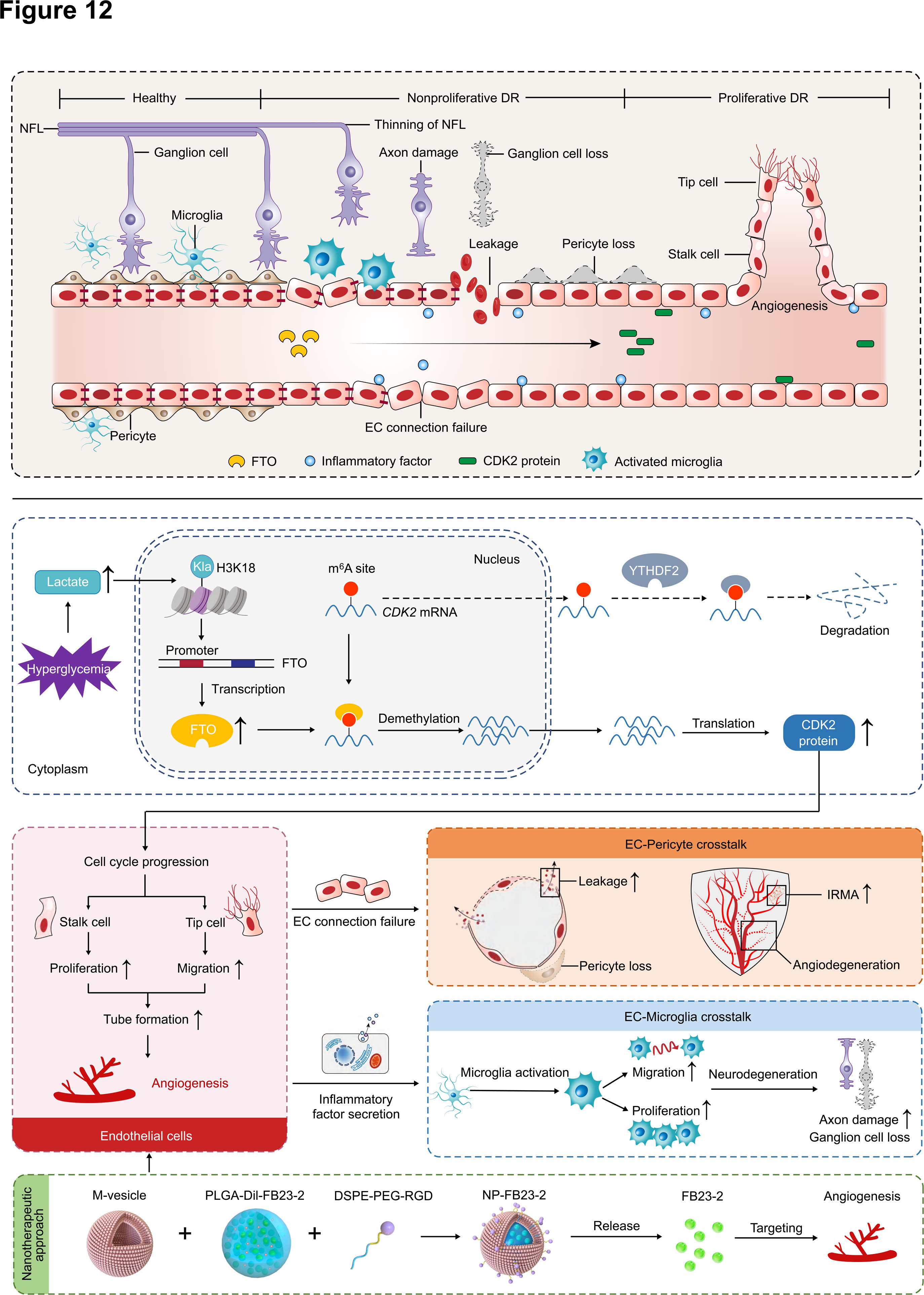
Schematic diagram of FTO-mediated effects on DR. Driven by histone lactylation H3K18la, FTO up-regulation in ECs triggers vascular endothelial tip cell formation and its crosstalk with pericyte and microglia to aggravate diabetes-induced microvascular dysfunction. FTO mediates DR phenotypes via regulating *CDK2* mRNA stability with YTHDF2 as the reader in an m^6^A-dependent manner.

Recent studies identified that dysregulated m^6^A modification mediated by aberrantly expressed m^6^A writers, erasers and readers contributes to DR progression, showing involvements in various pathological processes of DR. In ECs, Cao et al reported the DR promoting effects of METTL3 by regulating endothelial-mesenchymal transition via the SNHG7/KHSRP/MKL1 axis (45), while Zhao et al revealed its DR inhibitory role by mediating endothelial tight and adherens junctions in an ANXA1-dependent manner (46). YTHDF2 inhibits EC proliferation and tube formation to alleviate DR progression by mediating *ITGB1* mRNA instability (47). In perictytes, METTL3 governs pericyte dysfunction in DR by repressing PKC-η, FAT4 and PDGFRA expression through YTHDF2-dependent mRNA decay (10). In RPE cells, both METTL3 and YTHDF2 show alleviatory roles in high glucose induced RPE pyroptosis (44, 48). Additionally, dysregulated m^6^A modification also contributes to diabetes correlated retinal inflammation. In DR, YTHDF2 reduces the activity and inflammatory responses in retinal Müller cells (47), and the m^6^A eraser ALKBH5 mediates m^6^A modification of A20 to enhance M1 polarization of retinal microglia (11). These findings imply the critical and extensive involvements of m^6^A regulators in all pathological processes of DR. Nevertheless, the role of m^6^A eraser FTO in DR has been rarely discussed.

FTO dysregulation contributes to various oculopathies and participates in multiple pathogenesis. Reportedly, FTO promotes corneal neovascularization by regulating EC functions in an m^6^A-YTHDF2-dependent manner (49). FTO also alleviates AMD by preventing Aβ1-40 induced RPE degeneration via the PKA/CREB signaling pathway (50). Moreover, FTO shows uveitis curing effects by modulating m^6^A level of ATF4 to suppress inflammatory cytokine secretion and maintain tight junctions in RPE cells (51). Although FTO’s role in DR has not been annotated before, its involvement in diabetes and glucose metabolism has been widely discussed. Association between *FTO* gene polymorphisms and type 2 diabetes has been verified in many races (52, 53), supporting the clinical correlation between FTO and diabetes. Elevated *FTO* mRNA expression is detected in peripheral blood samples from type 2 diabetic patients, resulting in decreased m^6^A content and disturbed glucose metabolism (54, 55). Noteworthy, accumulating evidence suggests the direct regulation of FTO on glucose metabolism (56, 57). These findings provide fresh insights into the regulatory roles of FTO in oculopathies and diabetes. Herein, our study demonstrated the pathological involvement of FTO in DR. We revealed that FTO up-regulation in EC contributes to DR features by mediating *CDK2* expression in an m^6^A-YTHDF2-dependent manner. However, other potential downstream targets and pathways of FTO in ECs may also exist, which needs to be explored systematically.

Investigating the upstream transcriptional regulation of FTO in EC is important for better understanding into the molecular mechanism of DR. Guo et al elucidated that the transcription factor Foxa2 negatively regulates the basal transcription and expression of FTO in HEK293 cells (58). In mouse TM3 cells, aromatic hydrocarbon receptor affects FTO expression through transcriptional regulation (59). STAT3 also binds with FTO promoter to activate its transcription in breast cancer cells (60). Herein, we noticed that FTO up-regulation in EC under diabetic conditions is driven by lactate mediated histone lactylation H3K18la. Lactylation, which links metabolism and gene regulation, shows deep and extensive involvements in biological and pathological processes (39, 61, 62). Recent studies identify the regulation of lactylation on oculopathies, while its role in DR has not been elucidated. Lactylation of YY1 in microglia promotes retinal angiogenesis through transcription activation-mediated up-regulation of FGF2 (63). Histone lactylation was found to drive oncogenesis by facilitating YTHDF2 expression in ocular melanoma (61). In addition, lactylation also mediates METTL3 expression to promote immunosuppression of tumor-infiltrating myeloid cells (14), further emphasizing the interaction between lactylation and m^6^A modification. However, we could not completely exclude that other factors may be responsible for FTO up-regulation under diabetic conditions due to the complicated etiology and tangled process of DR. Thus, more comprehensive and in-depth investigations on initiating molecular events in DR that triggers FTO dysregulation are warranted.

Herein, we also developed and characterized a macrophage membrane coated and PLGA-Dil based nanoplatform loaded with the FTO inhibitor FB23-2 to treat DR. The macrophage membrane enhances the accumulation of nanoparticles in retinal angiogenic lesions, and the PLGA-based nanosystem ensures the controlled release of FB23-2 (41). Targeting efficacy of NP-FB23-2 was validated both *in vivo* and *in vitro*. NP-FB23-2s exhibited enhanced cellular uptake into ECs and achieved high concentration in retinal neovasculatures in mice receiving intravenous administration. Moreover, therapeutic efficiency of NP-FB23-2s on promoting m^6^A content and suppressing retinal neovascularization was also validated in mice, providing novel insights and options for therapeutic strategies of DR.

In conclusion, we demonstrated the pathological involvement of m^6^A demethylase FTO in DR. FTO overexpression triggered a series of diabetic retinal phenotypes, including angiogenesis, vascular leakage, inflammation and neurodegeneration, through directly affecting EC features and mediating EC-perictyte/microglia interactions. We also annotated the up- and down-stream regulatory network of FTO in ECs, and developed a novel macrophage membrane coated and PLGA-Dil based nanoplatform encapsulating the FTO inhibitor FB23-2 for systemic administration. This work describes an FTO-mediated regulatory network that coordinates EC biology and retinal homeostasis in DR, providing novel insights into DR pathogenesis, and indicates a promising nanotherapeutic approach for DR.

## Materials and Methods

### Ethical approval

Retinal proliferative membranes from DR patients and ERMs from age-matched controls without diabetes were obtained from Department of Ophthalmology in The First Affiliated Hospital of Nanjing Medical University. All procedures followed the Association for Research in Vision and Ophthalmology (ARVO) statement on human subjects and the Declaration of Helsinki with written informed consents signed by all individuals before donation. This study was approved and reviewed by the ethical committees from The First Affiliated Hospital of Nanjing Medical University (Approval No.: 2020-SR-545). Animal experiments, conformed to the guidelines of the Care and Use of Laboratory Animals (published by the NIH publication No. 86-23, revised 1996), were approved and consistently reviewed by the ethical review board of Nanjing Medical University (Approval No.: IACUC-2203035 for mice; IACUC-2204056 for zebrafish).

### Cell culture and treatment

HUVECs were cultured in DMEM/Low Glucose medium or DMEM/High Glucose medium supplemented with 10% fetal bovine serum (FBS; Invitrogen, Carlsbad, CA, USA), penicillin (100 U/mL; Invitrogen) and streptomycin (100 U/mL; Invitrogen). HMC3 cells were maintained in DMEM/F12 added with 10% FBS, penicillin (100 U/mL) and streptomycin (100 U/mL). RAW 264.7 cells were cultured in DMEM/High Glucose medium supplemented with 10% FBS, penicillin (100 U/mL) and streptomycin (100 U/mL). Complete medium was used short for supplemented culture medium in the following text. Cells were maintained at 37 °C with 21% O_2_ and 5% CO_2_. Co-cultivation of HUVECs and HMC3 cells was achieved using 0.4-μm-pore size Transwell chambers as indicated in **Figure 5D**. For actinomycin D assay, HUVECs were cultured in complete medium added with actinomycin D (1 µg/mL), and were harvested at, 4, 6, 8 and 12 hours post treatment, respectively. For lactate and C646 treatment, HUVECs were maintained in complete medium supplemented with L-lactate (10 mM; Sigma-Aldrich, St. Louis, MO, USA) or C646 (100 µM; MCE, Monmouth Junction, NJ, USA) for 48 hours before collection. For FB23-2 and NP-FB23-2 treatment, HUVECs were cultured in complete medium added with FB23-2 (2 µM; MedChemExpress, Princeton, NJ, USA) or NP-FB23-2 (dilution: 1:100) and harvested at 48 hours post treatment.

### RNA isolation and qCPR

Total RNA was obtained from cell lysates and mice retina using TRIzol reagent (Invitrogen). Nano-Drop ND-1000 spectrophotometer (Nano-Drop Technologies, Wilmington, DE, USA) was applied to detect the concentration and purity of RNA. cDNA was generated using a PrimeScript RT Kit (Takara, Otsu, Shiga, Japan). RNA levels were determined by qPCR with FastStart Universal SYBR Green Master (ROX; Roche, Basel, Switzerland) using StepOne Plus Real-Time PCR System (Applied Biosystems, Darmstadt, Germany). To normalize mRNA expression, GAPDH expression was assessed in parallel. Detailed primer information was listed in **Supplementary Table S1**.

### M^6^A dot blot assay

Extracted RNA was diluted, denatured, and loaded onto the Amersham Hybond-N+ membrane (Millipore, Boston, MA, USA). The membranes were then subjected to UV cross-linking (10 minutes), methylene blue (MB) staining (5 minutes) and 5% BSA blocking (1 hour). After that, the membranes were treated with m^6^A antibody at 4 °C overnight, and incubated with horseradish peroxidase (HRP)-conjugated secondary antibody (dilution: 1:5000; ICL, Newberg, OR, USA) at room temperature for an hour. The Tanon-5200Multi Chemiluminescent Imaging System (Tanon Science & Technology, Shanghai, China) was used to visualize the blots. Image J software (http://rsb.info.nih.gov/ij/index.html) was applied to measure dot intensity.

### M^6^A RNA methylation quantification assay

The m^6^A level of total RNA was detected using the EpiQuik m^6^A RNA Methylation Quantification Kit (Epigentek Group, Farmingdale, NY, USA) according to the manufacturer’s protocol. Briefly, 200 ng of extracted RNA, negative and positive controls were coated on assay wells, incubated with capture antibody solution and treated with detection antibody solution. The m^6^A level was then determined using a Synergy 4 automatic microplate reader (Agilent BioTek; Winooski, VT, USA) at 450 nm.

### Mouse breeding and manipulations

C57BL/6J mice were raised in a specific pathogen-free facility in Nanjing Medical University with a 12h light/dark cycle at 28.5 °C. Embryos were produced through natural mating. No randomization was applied. Before all invasive operations and examinations, mice were fully anesthetized through intra-peritoneal injection of ketamine (80 mg/kg) and xylazine (4 mg/kg) with pupils dilated using 1% cyclopentolate-HCL and 2.5% phenylephrine.

STZ mice were generated by continuous intra-peritoneal injection of STZ (50 mg/kg; Sigma-Aldrich) for 5 days. Before injection, 6-week-old male mice were in abrosia for 8 hours. Water was supplied during fasting. Blood glucose was continuously measured till 7 days after injection using a Contour TS blood glucose monitor glucometer (Bayer, Leverkusen, Germany). Mice with blood glucose of over 15 mmol/L were considered as diabetic mice and were selected for further investigations. To generate OIR mice, P7 neonatal mice together with their nursing mother were housed in a closed chamber with 75% O_2_ for 5 days. Food and water were supplied as normal.

For intra-vitreal injection, 1 µL solution containing AAV-blank/AAV-Fto supernatant (10^12^∼10^13^ genome copies/mL; AAV Serotype 2) was delivered into the vitreous chamber of mice using a syringe with a 33-gauge needle (Hamilton, Bonaduz, Switzerland). For intra-cardiac injection of Texas red dextran, mouse thoracic cavity was carefully opened to expose the heart. One mL Texas red dextran solution (2 mg/mL; Invitrogen) was injected into the left ventricle at an even speed for 1 minute. Mouse was sacrificed immediately after perfusion with eyes enucleated to produce retinal flat mounts. For intravenous injection, Evans blue solution (4 mL/kg; Sigma-Aldrich) was administered through the tail vein. Mice were sacrificed with eyes enucleated at 2 hours post injection to make retinal flat mounts. NP-FB23-2 (3 mg/kg) was delivered through the retro-orbital sinus into neonatal mice with a 31-gauge insulin syringe (BD Biosciences, San Jose, CA, USA).

### Mounting of mice retina

For tissue preparation, mice were anesthetized and sacrificed after the ophthalmic examinations with eyes enucleated and connective tissues trimmed. After careful removal of the anterior segments and vitreous, the remaining posterior eyecups were used for further tests. For dot blot, qPCR and immunoblotting assays, neural retinas were isolated and sent for RNA and protein extractions. For retinal flat mounting staining, eyecups were fixed in 4% paraformaldehyde (PFA) at 4 °C for at least 2 hours. Neural retinas were then carefully separated from the posterior eyecup and trimmed into a four-leaf clover-shaped retinal flat mount.

### Immunoblotting

For protein isolation, cells and mice tissue were collected and fragmented in lysis buffer (Beyotime, Shanghai, China) supplemented with protease inhibitors cocktail (Roche). Isolated proteins were then segregated by gel electrophoresis and transferred to polyvinylidene fluoride membranes (Millipore). Membranes were subsequently incubated in primary antibodies (**Supplementary Table S2**) at 4 °C overnight and probed by HRP-conjugated secondary antibodies (dilution: 1:10000; ICL) for an hour at room temperature. Blots were developed using the Tanon-5200Multi Chemiluminescent Imaging System. Proteins were quantified using Image J software.

### Cell transduction and transfection

Empty lentiviral plasmid with FLAG tag (Ubi-MCS-3FLAG-SV40-puromycin; L-EV), lentiviral plasmid containing the CDS of wild type human *FTO* gene (Ubi-MCS-FTO-3FLAG-SV40-puromycin; L-FTO) and with the two catalytically inactive mutations H231A and D233A (Ubi-MCS-FTO^MU-^3FLAG-SV40-puromycin; L-FTO^MU^) were constructed by GeneChem (Shanghai, China). Lentiviral plasmid L-EV, L-FTO, or L-FTO^MU^ was co-transfected with pCMV-Gag/Pol and pCMV-VSVG into HEK293T cells using Lipofectamine 3000 transfection reagent (Invitrogen) according to the manufacturer’s instructions. Packaged lentiviral particles in the supernatant were collected at 48 hours post-transfection, mixed with polybrene (2 µg/mL; Sigma-Aldrich), and added into HUVECs at a multiplicity of infection (MOI) of 10. Transduced HUVECs were sorted out using puromycin (5 µg/mL; Sigma-Aldrich) and sent for further analyses. For transfection assay, HUVECs were transfected with indicated siRNAs using Lipofectamine 3000. Scramble siRNA and siRNAs targeting *FTO* or *YTHDF2* were purchased from RiboBio (Guangzhou, China) with sequences detailed in **Supplementary Table S3**.

### Immunofluorescence staining

Retinal proliferative membranes, ERMs, cells, and posterior eyecups of mice were fixed in 4% PFA. Neural retinas were separated from the posterior eyecups to obtain neural retinal flat mounts. After being permeabilized with 0.5% Triton X-100 (Sigma-Aldrich) and blocked in 1% BSA, retinal proliferative membranes, ERMs, cells and neural retinal flat mounts were sequentially incubated with primary antibodies (**Supplementary Table S2**) at 4 °C overnight, and corresponding fluorescence-conjugated secondary antibodies (dilution: 1:100 for tissue staining; 1:1000 for cell staining; Invitrogen) at room temperature for 2 hours. Cell nuclei were counterstained by DAPI (Sigma-Aldrich). Fluorescence was observed with an upright microscope (DM4000 B, Leica), a cell imaging multimode reader (BioTek Cytation 1, Agilent BioTek) and an inverted microscope (DMi8, Leica) equipped with THUNDER imaging system (Leica). Fluorescence intensity was quantified with Image J software.

### RNA-Seq, MeRIP-Seq and MeRIP-qPCR

Extracted total RNAs with high quality was sent for rRNA depletion using a human TransNGS rRNA Depletion Kit (TransGen Biotech, Beijing, China) and RNA purification with MagicPure RNA beads (TransGen Biotech). RNAs were then chemically segmented into 60∼200 nt fragmentations with 10% of total RNA separated as input control (RNA-Seq). The rest RNAs were incubated with Dynabeads Protein A (Thermo Fisher Scientific, Waltham, MA, USA) and m^6^A antibody at 4 °C for 2 hours. M^6^A RNA enrichment was then detected via high-throughput sequencing (MeRIP-Seq) or qPCR (MeRIP-qPCR). For RNA-Seq and MeRIP-Seq, purified input RNAs and immunoprecipitated RNA fragments were sent for library construction with the VAHTS Total RNA-Seq (H/M/R) Library Prep Kit for Illumina (Vazyme, Nanjing, China) and sequencing using the Illumina HiSeq 2500 (Illumina, San Diego, CA, USA). Quality control and raw sequencing data filtration were achieved with the FastQC and fastp software. Reads were then aligned to the human genome GRCh38/hg38 using STAR software. Reads per kilobase per million mapped reads (RPKM) values were measured in 5’-UTR, CDS, and 3’-UTR of all genes using deepTools software. For RNA-Seq, expression of mRNAs in input RNAs was analyzed using Stringtie algorithm with differential expression annotated by Deseq2 software. For MeRIP-Seq, Methylated MeRIP peaks were called using exomePeak2 software, annotated with Annovar software, and matched to motifs using Homer software. For MeRIP-qPCR, immunoprecipitated RNA fragments as well as input RNAs were reversely transcribed into cDNAs. Enrichment of m^6^A-immunopurified *FTO* mRNA was measured with qPCR and normalized to the input.

### Cell cycle analyses

Cell cycle was analyzed using the cell cycle staining kit (Multisciences Biotech, Hangzhou, China) according to the manufacturer’s instructions. Collected cells were suspended in DNA staining solution containing permeabilization solution for 30 minutes. Cell cycle was the detected by flow cytometry (Beckman Coulter, Brea, CA, USA) with data analyzed using the win cycle software. A total of 10000 cells from each sample were measured to modulate the cell cycle.

### EdU assay

Cell proliferation was determined using the EdU Apollo In Vitro Kit (RiboBio) per the manufacturer’s protocols. Cells were incubated in EdU solution for 2 hours, fixed with 4% PFA, permeabilized using 0.5% Triton X-100, and fluorescently labeled in Apollo solution. Cell nuclei were counterstained with DAPI. Proliferating cells showing red signals were visualized using a DM4000 B upright microscope or an inverted microscope (DMi8) equipped with THUNDER imaging system.

### Transwell migration assay

Cell migration was examined using an 8-μm-pore size Transwell migration chamber. Cells were seed into the upper chamber with its bottom immersed in culture medium. Cell migration was allowed to proceed for 24 hours at 37 °C with 21% O_2_ and 5% CO_2_ before collection. Cells that migrated to the below surface of the upper chamber were then stained with crystal violet (Beyotime) for 15 minutes. Different views were randomly chosen and recorded with an ECLIPSE Ts2 inverted microscope (Nikon, Tokyo, Japan) with average counting taken.

### Scratch test

Cell migration was also determined using scratch test. HUVEC monolayers were scraped in straight lines with pipet tips to create scratches, and gently washed to remove cell debris and smooth scratch edges. Images of the same scratch site were recorded right after the scratch and at 24 hours after the scratch using an ECLIPSE Ts2 inverted microscope, and were analyzed with the Image J software.

### Migration assay with the RTCA system

Real-time rates of HUVECs migration were detected using the RTCA system (Roche) according to the manufacturer’s protocol. Briefly, cells were seeded into the E-Plate and maintained in complete medium. Impedance value for each well was automatically recorded by the RTCA system as a CI value. Migration rates were calculated based on the slope of the line between two given time points.

### Tube formation assay

HUVECs were seeded onto growth factor-reduced matrigel (BD Biosciences) in 24-well plates to observe formation of capillary-like structures. Images were taken at 5 hours post plantation using an ECLIPSE Ts2 inverted microscope. Image J software was applied to quantify node number and branching length of HUVECs.

### mRNA synthesis

CDS of zebrafish *fto* mRNA was synthesized and inserted into the pCS2+ plasmid (Addgene, Watertown, MA, USA) to get the linearized pCS2+-FTO plasmid for *in vivo* transcription. The inserted sequence of the recombinant plasmid was validated using Sanger sequencing in both directions. Capped and tailed zebrafish *fto* mRNA was generated with a mMESSAGE mMACHINE T7 Ultra Kit (Ambion, Austin, TX, USA) and purified using an RNeasy Mini Kit (Qiagen, Hilden, Germany) according to the manufacturers’ protocol.

### Zebrafish manipulations

The transgenic zebrafish strain *Tg(LR57:GFP)* was a kind gift from the Zebrafish Center of Nantong University. One- to two-cell-stage [(0 hour post fertilization (hpf)] zebrafish embryos were collected and randomly microinjected with one nano-liter (nL) solution containing different dosages of purified *fto* mRNA or antisense mRNA. At 24 hpf, embryos with systemic deformities were discarded. The rest embryos were further randomly divided into two groups and maintained in fish-raising water with or without 3% glucose. At 48 hpf, embryos receiving distinct treatments and showing normal systemic appearance were selected for vascular imaging. Truncal and ocular vasculatures of living zebrafish were visualized using a Nikon A1 HD25 confocal microscope system (Nikon).

### Ophthalmic examinations

For fundus examinations, mice were anesthetized with pupils dilated. Fundus photo was taken using a ZEISS CLARUS 500 fundus camera (Carl Zeiss, Jena, Germany). FFA was conducted after intraperitoneal injection of fluorescein sodium (International Medication Systems, South El Monte, CA, USA) at 2 µL/g body weight. Fluorescent fundus images were acquired through Heidelberg Retina Angiograph 2 (Heidelberg Engineering, Heidelberg, Germany).

### Retinal trypsin digestion and PAS staining

Fixed and isolated neural retinas were washed in distilled water overnight and digested in 3% trypsin (BioFroxx GmbH, Einhausen, Germany) at 37 °C for an hour. Digested retinas were carefully washed and gently shaken with a pipette under the microscope to separate the retinal vasculatures. For PAS staining, the isolated retinal vessels were sequentially stained with periodic acid solution and Schiff’s reagent. Images were taken with an upright microscope (DM4000 B).

### TMT-based quantitative proteomic analysis

Cells were harvested with intra-cellular proteins extracted and digested with trypsin. Peptides were then labeled with TMT using a TMT10plex mass tag labeling kit (Thermo Fisher Scientific) per the manufacturer’s instructions. TMT-labeled peptides were then separated by high pH reverse-phase high performance liquid chromatography (HPLC) with C18 columns (Agilent BioTek) and dried in a vacuum centrifuge. Liquid chromatography-tandem mass spectrometry (LC-MS/MS) was subsequently performed. All fragments were sequentially dissolved in aqueous solution (0.1% formic acid and 2% acetonitrile), loaded onto a home-made reverse-phase analytical column, and subjected to an EASY-nLC™ 1000 ultraperformance liquid chromatography (UPLC) system (Thermo Fisher Scientific) at a constant flow rate of 400 nL/minuets. For MS settings, the applied electrospray voltage was 2.0 kV and the m/z range was 350 to 1800 for the complete scan. Peptide fragments were quantified with Parallel Reaction Monitoring (PRM). Proteins were identified by Swissprot database and quantified using Proteome Discoverer 2.0.

### LiquiChip

Culture medium of HUVECs were collected and analyzed with the Bio-Plex Pro™ Human Cytokine 27-plex Panel (Bio-Rad Laboratories, Hercules, CA, USA) in accordance with the manufacturer’s protocols. Briefly, culture medium was sequentially mixed with fluorescence-labeled capture beads and highly specific monoclonal antibodies. The mixture was then resuspended and irradiated. Concentrations of 27 cytokines and chemokines in the mixture were detected using the Luminex™ 200™ instrument system (Luminex, Austin, TX, USA). Data were analyzed with the xPONENT® software.

### RIP-qPCR

RIP experiment was performed using the Magna RIP™ Kit (Millipore) per the manufacturer’s instructions. Protein A/G magnetic beads were incubated with antibodies against IgG, FTO and YTHDF2 respectively. qPCR was used to measure the enrichment of protein-bounded RNA. Results were normalized to the input.

### Lactate content analysis

Lactate concentration was determined using a Lactic Acid Content Assay Kit (Solarbio, Beijing, China) according to the manufacturer’s protocol. HUVECs were harvested, sonicated and incubated with indicated reagents. Cell deposits were extracted and dissolved in absolute alcohol. Lactate content was monitored using a Synergy 4 automatic microplate reader (Agilent BioTek) at 570 nm.

### ChIP-qPCR

ChIP assay was conducted using the SimpleChIP^®^ Enzymatic Chromatin IP Kit (Cell Signaling Technology, Danvers, Massachusetts, USA) per the manufacturer’s instructions. Cells were crosslinked with 1% formaldehyde, sonicated and digested to acquire chromatin fragments. DNA fragments were collected and incubated with H3K18la antibody at 4 °C overnight. Enrichment of H3K18la-bounded DNA was measured using qPCR and normalized to the input.

### Preparation of NP-FB23-2

We used nanoprecipitation method to prepare PLGA nanoparticles encapsulating FB23-2. Briefly, PLGA (5 mg, MW=100,000; Daigang Biology, Jinan, China), distearoyl phosphoethanolamine-poly (DSPE-PEG-COOH; 2 mg; Qiyue Biology, Xi’an, China), 1, 2-Dioleoyl-sn-glycero-3-phosphocholine (DOPC; 1 mg; Qiyue Biology) and FB23-2 (2 mg) were dissolved in 1 mL organic phase. The mixture was added dropwise into 4 mL PBS buffer with stirring. The suspension was then sonicated at 100W for 15 minutes and dialyzed in a 3500DA dialysis bag for 24 hours. Next, 0.1 wt% DiI (Beyotime) was loaded into the collected PLGA-FB23-2 solution to obtain fluorescently labeled PLGA-Dil-FB23-2 nanoparticles.

To obtain purified macrophage membranes, RAW 264.7 cells were collected and treated according to a previously described protocol (41). Cells were resuspended in ice-cold 0.1×TM buffer (Beyotime) containing protease and phosphatase inhibitor cocktail (Thermo Fisher Scientific), and disrupted by freezing, thawing and a homogenizer (IKA, Staufen, Germany). The suspension was then sent for three rounds of centrifugations (3200g for 5 minutes; 20,000g for 25 minutes; 100,000g for 60 minutes) to get M-vesicles. The PLGA-Dil-FB23-2 nanoparticles and the M-vesicles were sonicated separately for 5 minutes and subsequently mixed at a 1:1 (w/w) ratio of membrane protein to polymer. The mixture was sequentially extruded through 400 nm and 200 nm polycarbonate membranes using a micro-extruder (Avestin, Ottawa, Canada) for 10 rounds to obtain macrophage membrane coated PLGA-Dil-FB23-2s nanoparticles.

Finally, DSPE-PEG-RGD (1 mg; Qiyue Biology) dissolved in 0.5 mL of N, N-Dimethylformamide (DMF; Sigma-Aldrich) was added to the macrophage membrane coated PLGA-Dil-FB23-2s nanoparticles under stirring. The nanoparticle solution was then dialyzed in a 10,000DA dialysis bag for 24 hours, and concentrated using a 10,000DA MWCO ultra centrifugal filter tube (Millipore) to obtain NP-FB23-2.

### TEM and DLS

To visualize NP-FB23-2, solution containing NP-FB23-2 was dropped onto a copper and observed using an FEI Tecnai G2 Spirit BioTWIN electron microscope (FEI, Hillsboro, OR, USA). Size of NP-FB23-2 was measured using the NanoBrook 90Plus particle size analyzer (Brookhaven Instruments, New York, NY, USA) based on the principles of DLS.

### H&E staining

To obtain paraffin sections of mice heart, liver, spleen, lung and kidney, tissues were isolated and fixed in 4% PFA, dehydrated and transparentized with conventional grade of alcohol and xylo, embedded in paraffin, and sectioned at 3∼5 µm thickness. Paraffin sections of mice tissues were dewaxed using xylo and conventional grade of alcohol, and subsequently stained with hematoxylin and eosin to visualize histological structures under a DM4000 B upright microscope.

### Quantification and statistical analysis

GraphPad Prism (v 4.0; GraphPad Software, San Diego, CA, USA) was used for statistical analyses. Two tailed student’s T test was utilized for comparison between two groups. One-way analysis of variance (ANOVA) coupled with the Bonferroni’s post hoc test was used for comparisons among three groups. Data were presented as mean ± standard error of the mean (SEM). P < 0.05 was considered of statistical significance. Detailed replicate information for each experiment was written in the figure legends.

## Acknowledgments

We thank all donors for their donation. We are grateful to Prof Dong Li and Dr Xin-Yu Wang (Institute of Biophysics, Chinese Academy of Sciences) for technical supports on MeRIP-Seq, and Prof Dong Liu (Nantong University) for supports on zebrafish experiments. This study was supported by National Natural Science Foundation of China (82070974 to XC, 82271100 and 81970821 to QHL); Jiangsu “333” Advanced Talent-training Project to XC. The funders had no role in study design, data collection and analysis, decision to publish, or preparation of the manuscript.

## Author contributions

XC, RXS and JNW contributed equally to this work. XC designed research studies. RXS, JNW, YRZ, QB, YCZ, HJZ and JXZ conducted experiments. XC, RXS, JNW, YRZ, QB and YCZ acquired and analyzed data. WWZ, JDJ and STY provided intellectual input or provided reagents. XC and QHL conceived the study and secured funding. QHL supervised the work. XC, RXS and JNW wrote the manuscript with support from all authors. The order of co-first authors was decided by discussion among the three first authors and the corresponding author.

